# GDNF receptor agonist supports dopamine neurons in vitro and protects their function in animal model of Parkinson’s

**DOI:** 10.1101/540021

**Authors:** Arun Kumar Mahato, Juho-Matti Renko, Jaakko Kopra, Tanel Visnapuu, Ilari Korhonen, Nita Pulkkinen, Maxim Bespalov, Eric Ronken, T. Petteri Piepponen, Merja Voutilainen, Raimo K. Tuominen, Mati Karelson, Yulia A. Sidorova, Mart Saarma

## Abstract

Motor symptoms of Parkinson’s disease (PD) are caused by degeneration and progressive loss of nigrostriatal dopamine neurons. Currently no cure for this disease is available. Existing drugs alleviate PD symptoms, but fail to halt neurodegeneration. Glial cell line-derived neurotrophic factor (GDNF) is able to protect and repair dopamine neurons *in vitro* and in animal models of PD, but its clinical use is complicated by pharmacokinetic properties. In the present study we demonstrate the ability of a small molecule agonist of GDNF receptor RET to support the survival of cultured dopamine neurons only when they express GDNF receptors. In addition, BT13 activates intracellular signaling cascades *in vivo*, stimulates release of dopamine and protect the function of dopaminergic neurons in a 6-hydroxydopamine (6-OHDA) rat model of PD. In contrast to GDNF, BT13 is able to penetrate through the blood-brain-barrier. Thus, BT13 serves as an excellent tool compound for the development of novel disease-modifying treatments against PD.

## Introduction

Parkinson’s disease is the second most common neurodegenerative disease, affecting 1–2% of the population over 50 years of age with a higher frequency in males (Samii *et al.*, 2004; Thomas and Beal, 2007). The characteristic motor symptoms of Parkinson’s disease, including bradykinesia, rigidity, resting tremor, postural and gait impairment, result from the progressive degeneration and death of dopamine neurons in the substantia nigra pars compacta (SNpc) rendering dopamine deficiency within the basal ganglia. (Dauer and Przedborski, 2003; Kalia and Lang, 2015). Patients with Parkinson’s disease also suffer from multiple non-motor symptoms (Kalia and Lang, 2015; Lee and Koh, 2015; Schapira *et al.*, 2017). Current drugs for Parkinson’s disease management are either supplementing or mimicking endogenous dopamine. While these treatments provide symptomatic relief, they neither halt nor reverse the disease progression and have very little effect on non-motor symptoms. Therefore, the major challenge remains to develop treatments that protect and restore dopamine neurons in Parkinson’s disease patients and alleviate non-motor symptoms.

Glial cell line-derived neurotrophic factor (GDNF) and the related protein neurturin (NRTN), belonging to the GDNF family ligands (GFLs), have been identified and characterized for their ability to promote the survival of cultured nigrostriatal dopamine neurons and to protect and repair them in animal models of Parkinson’s disease (Lin *et al.*, 1993; Hoffer *et al.*, 1994; Kojima *et al.*, 1997; Horger *et al.*, 1998; Kirik *et al.*, 2000; Li *et al.*, 2003; Kordower *et al.*, 2006). That is why GDNF and NRTN were also tested in patients with Parkinson’s disease. Despite promising results in open-label Phase I/II clinical trials with both GDNF protein (Gill *et al.*, 2003; Slevin *et al.*, 2005) and adeno-associated virus vector-encoded NRTN (AAV2-NRTN) (Marks *et al.*, 2008), randomized placebo-controlled Phase II clinical trials failed to reach primary end-points (Nutt *et al.*, 2003; Lang *et al.*, 2006; Marks *et al.*, 2010; Olanow *et al.*, 2015). It should be noted, however, that early-stage Parkinson’s disease patients, in contrast to late-stage patients, might respond to AAV2-NRTN treatment (Bartus and Johnson, 2017). Thus, phase III clinical trial with GDNF protein in patients with Parkinson’s disease of moderate severity is currently under discussion with regulatory agencies (http://medgenesis.com/news.htm#Advice).

The poor pharmacokinetic properties of GFL proteins, necessity for intracranial delivery via stereotaxic surgery, inconsistent biological activity and high price make GFLs themselves poor drug candidates for the treatment of Parkinson’s disease. However, small molecule synthetic compounds with blood-brain barrier (BBB) penetrating properties targeting GFL receptors and mimicking GFLs biological effects in dopamine neurons may translate to clinic easier. Such compounds can affect both terminal regions of dopamine neurons inducing sprouting and their cell bodies promoting survival, thus producing superior effects in comparison to GDNF. Systemically delivered GFL mimetics may also alleviate the non-motor symptoms of Parkinson’s disease often caused by degeneration or dysfunction of GFL-responsive brain and peripheral neurons (Pichel *et al.*, 1996; Paratcha *et al.*, 2006).

GFLs signal via receptor tyrosine kinase RET. GDNF and NTRN first bind to glycosylphosphatidylinositol-anchored GDNF family receptor α1 and α2 (GFRα1 and GFRα2), respectively, which then form a complex with RET and stimulate RET autophosphorylation. This leads to activation of intracellular signaling cascades important for neuronal survival and function (Airaksinen and Saarma, 2002; Ibáñez, 2013).

Previously, we discovered a small molecule, BT13, that selectively activates GFL receptor-dependent signaling in immortalized cells, supports sensory neurons and alleviates neuropathy in rats (Sidorova et al., 2017). In the present study, we evaluated the biological effects of BT13 in dopamine system. We confirmed that BT13 activates RET and downstream signaling cascades in cells expressing GDNF and NRTN receptors. We found that 0.1–1 μM BT13 supports the survival of cultured dopamine neurons from wild-type, but not from RET knock-out embryos. In the striata of intact mice, infusion of BT13 stimulated dopamine release as determined with microdialysis. Finally, BT13 reduced amphetamine-induced rotational behavior in 6-hydroxydopamine (6-OHDA) rat model of Parkinson’s disease. Therefore, BT13 is considered as a valuable tool compound for the development of a novel treatment for Parkinson’s disease.

## Materials and methods

Experimental details are presented in Expanded View section.

### Cell Lines

MG87RET cells (Eketjäll *et al.*, 1999) and reporter cell lines on their basis described previously (Sidorova *et al.*, 2010).

### Plasmids

Full-length human GFRα1 cDNA subcloned in pCDNA6 (Invitrogen, USA) (Sidorova *et al.*, 2010), GFRα2 cDNA in pCR3.1 and enhanced green fluorescent protein (GFP) cDNA in pEGFP-N1.

### Proteins

Human recombinant GDNF for *in vitro* experiments from Icosagen (Estonia), for in vivo studies - from PeproTech (USA).

### BT13

BT13 was synthesized by EvoBlocks Ltd (Hungary) and had 98.6% purity (LCMS). The structure, integrity, and purity of BT13 was verified by NMR experiments recorded on a Bruker Avance III HD NMR spectrometer operated at ^1^H frequency of 850.4 MHz equipped with a cryogenic probe head by NMR facility at the Institute of Biotechnology, University of Helsinki. BT13 synthesis scheme and NMR data are provided in Expanded View section.

### Experimental animals

Adult male mice C57Bl/6 (JRccHsd) (Harlan, the Netherlands) weighing 19–32 grams (8-15 weeks) were used for brain microdialysis experiments and for immunohistochemistry of phosphorylated extracellular-signal-regulated kinases 1 and 2 (pERK) and phosphorylated ribosomal protein S6 (pS6) in striatum. Primary dopamine neurons were dissected from E13.5 embryos of NMRI and RET knock-out mice (C57BL/6JOlaHsd) (Laboratory Animal Centre, University of Helsinki). Adult male Wistar rats (RccHan:WIST) (Harlan), weighing 240–435 grams at the start of the experiments were used for testing BT13 in the 6-OHDA model of Parkinson’s disease. All experiments were carried out according to the European Community guidelines for the use of experimental animals and approved by the National Animal Experiment Board of Finland (license numbers: ESAVI/7551/04.10.07/2013, ESAVI/198/04.10.07/2014) for experiments with living animals and the Laboratory Animal Centre of the University of Helsinki (license number: KEK15-022) for collection of E13.5 embryos of NMRI mice.

### Genotyping

Mouse E13.5 embryos were genotyped for the presence or absence of RET as described previously (Schuchardt *et al.*, 1996).

### Luciferase assay

Luciferase assay was performed as described previously (Sidorova *et al.*, 2010, 2017).

### Sample preparation for phosphorylation assays

Cells lysates for analysis of RET, pERK and pAKT phosphorylation in response to BT13 were prepared from MG87RET cells transfected with hGFRα1, hGFRα2 and GFP-expressing plasmids as described previously (Sidorova *et al.*, 2017).

### Western blotting-based ERK and AKT phosphorylation assay

Levels of pERK and pAKT in immortalized cells were analyzed by Western blotting using E4 pERK (1:1000, Santa Cruz Biotechnology Cat# sc-7383, RRID:AB_627545) and pAKT (1:500, Cell Signaling Technology Cat# 9271, RRID:AB_329825) antibodies as described previously (Sidorova *et al.*, 2017). Membranes were probed with GAPDH antibody (1:4000, Millipore Cat# MAB374, RRID:AB_2107445) to confirm equal loading.

### Analysis of RET phosphorylation by Western blotting and RET-ELISA

The level of RET phosphorylation in the cells was analyzed by Western blotting of immunoprecipitated RET with phospho-tyrosine-specific antibody (1:1500 Merck Millipore Cat# 05-321, RRID:AB_309678) as described previously (Leppanen *et al.*, 2004) or by RET-ELISA (Parkash *et al.*, 2008). Anti-RET C-20 antibody (Santa Cruz Biotechnology Cat# sc-1290, RRID:AB_631316) were used to confirm equal loading in Western blotting (1:500) and capture RET in RET-ELISA.

### Survival assay for naïve dopamine neurons

The effect of BT13 on survival of näive dopamine neurons was assessed using the method described previously (Planken *et al.*, 2010; Saarenpää *et al.*, 2017). Dopamine neurons were labelled with an antibody against the key enzyme of dopamine synthesis, tyrosine hydroxylase (TH) (1:500, Millipore Cat# MAB318, RRID:AB_2201528), on 5^th^ day in vitro (5^th^ DIV) followed by secondary antibody conjugated with fluorophore. Cells were imaged by CellInsight (CX51110, ThermoFisher Scientific) and analysed with CellProfiler image analysis software (Carpenter *et al.*, 2006).

### Survival of MPP^+^ challenged dopamine neurons

The survival analysis of MPP+ challenged wild-type dopamine neurons was performed by Neuron Experts company (http://www.neuronexperts.com/) using previously described method (Schinelli *et al.*, 1988). Cells were isolated from E15 rat brains and cultured for 6 days. On 6^th^ DIV cells were treated with 16 μM MPP^+^ and tested substances or DMSO as a negative control. The known survival promoting factor for dopamine neurons, brain-derived neurotrophic factor (BDNF) (10 ng/ml) (Hyman *et al.*, 1991), was used as a positive control. On 8^th^ DIV cells were fixed with 4% PFA and probed with antibody against TH (Sigma-Aldrich Cat# T1299, RRID:AB_477560) followed by fluorophore-conjugated secondary antibody. Cells were imaged using InCell AnalyzerTM 1000 (Amersham Biosciences, UK) and analyzed with InCell AnalyzerTM 1000 3.2. Workstation software.

### *In vivo* microdialysis

*In vivo* microdialysis was performed as described previously (Kumar *et al.*, 2015). BT13 was dissolved in Ringer solution until desirable concentration (the highest tested concentration (50 μM) was limited by BT13 solubility). The solutions were sonicated to enhance solubility of BT13. Vehicle (0.5% of DMSO) did not influence extracellular dopamine levels (data not shown). Concentration of dopamine was analyzed with high-performance liquid chromatography (HPLC). After achieving a stable baseline with Ringer solution, BT13 was delivered into the striatum as a continuous infusion by reverse dialysis until the end of the experiment.

### Microinjections and stereotaxic surgery

Bilateral microinjections into the mouse dorsal striatum were performed using stereotaxic frame (Stoelting, USA) using similar procedure as for *in vivo* microdialysis (Kumar *et al.*, 2015). Left striatum received BT13 (103.5 μg (≈100 μg) (N = 4), 207 μg (≈200 μg) (N = 4), 517.5 μg (≈500 μg) (N = 4) and 776.25 μg (≈750 μg) (N = 4)) or GDNF (5 or 10 μg (N = 4)) in saline with 0.5 % DMSO (vehicle). The right striatum was always injected with the vehicle. The animals were allowed to recover in their home cage on a heating pad for 1 hour after the injection. Afterwards the brains were collected for the analysis of pERK and pAKT levels.

### 6-Hydroxydopamine lesion

Catecholaminergic neurotoxin 6-OHDA (16 μg, Sigma-Aldrich, Germany; calculated as free base and dissolved in ice-cold saline with 0.02% ascorbic acid) was injected unilaterally into the left dorsal striatum as described previously (Voutilainen *et al.*, 2009). Desipramine (15mg/kg i.p.; calculated as free base; Sigma-Aldrich) was administrated 30 minutes before the 6-OHDA injection.

### Administration of BT13 and GDNF

BT13 (0.25-0.5 μg/μl in 100% propylene glycol), GDNF (0.25 μg/μl in PBS) or 100% propylene glycol (vehicle-treated group) were constantly infused into the striatum at a flow rate of ∼0.5 μl/h for 7 days starting 1 hour after the 6-OHDA injection via brain infusion cannula connected with Alzet minipump implanted as described earlier (Voutilainen *et al.*, 2011). The resulting dose for BT13 was ∼3-6 μg/24 h and for GDNF ∼3 μg/24 h. The dose for BT13 could not be determined accurately because the solubility of BT13 in 100% propylene glycol was limited to 0.5 μg/μl which was the highest concentration that remained dissolved for 7 days in stabile conditions in a test tube at 37°C. However, changes in ambient conditions during the pump implantation and 7-day infusion period may have precipitated some of the compound. In addition, variation in the mean pumping rate between different batches of the osmotic pumps may have resulted in dose deviation of more than 1 μg/24 h at the concentration of 0.5 μg/μl. Upon completion of the infusion, the pump, cannula, dental cement and screws were removed and the incision was sutured.

### Behavioral analysis

D-Amphetamine-induced rotational behavior was measured 2, 4 and 6 weeks after the 6-OHDA lesion in automatic rotometer bowls (Med Associates Inc., VT, USA) as described previously (Ungerstedt and Arbuthnott, 1970; Lindholm *et al.*, 2007). After completion of the last behavioral test, the brains were collected for analysis of dopamine system.

### Immunohistochemistry

Immunohistochemical stainings of pERK1/2 and pS6 were performed on 5 μm thick coronal sections of paraffin-embedded mouse brains using primary antibodies raised against pERK1/2 (1:300, Cell Signaling Technology Cat# 4370, RRID:AB_2315112) and pS6 (1:300, Cell Signaling Technology Cat# 5364, RRID:AB_10694233).

To assess the number of remaining dopamine neurons in the SNpc and the density of dopaminergic fibers in the striatum, free-floating 40-μm thick coronal cryosections were probed with antibodies against dopamine transporter (DAT) (1:2000, Millipore Cat# MAB369, RRID:AB_2190413) or TH (1:2000, Millipore Cat# MAB318, RRID:AB_2201528) as described previously (Voutilainen et al., 2009; Bäck et al., 2013). All stainings were visualized using respective secondary antibodies conjugated with HRP or ABC-system (Vectastain Elite ABC HRP Kit, Vector Laboratories Cat# PK-6100) and 3,3’-diaminobenzidine (DAB; Cat# SK-4100, USA) as a chromogen. Digital images of the striatal immunostained sections were acquired with an automated bright field microscopy slide scanner (3DHistech Ltd., Hunagry)

### Analysis of pERK and pS6 immunohistochemistry

Mean optical density (OD) of the staining in a single section (closest to the injection site and with the highest signal) from both hemispheres of each animal was analyzed using ImageJ software (Media Cybernetics Inc, USA). In cases where the highest signal was not located in the same section for both hemispheres, the values were counted from separate sections. OD values were counted in roughly equal sized areas from both hemispheres. Background OD values were measured from the peripheral striatal area that lacked the signal or from the septum and subtracted from the OD values in the areas selected for analysis. Resulting data were normalized to the area of analysed selections and subjected to the statistical analysis. For presentation purpose the values for BT13- and GDNF-treated sides were normalized to the values for vehicle-treated side of the same brain.

### Stereological assessment of TH-positive cells in the SNpc

TH-positive cell bodies in the SNpc of rat brain were counted by a person blinded to the treatment groups in the SNpc on 3 coronal sections at approximately the same rostro-caudal sites using Olympus BX51 (Olympus Corporation, Japan) and Stereo Investigator software version 11.06.2 (MBF Bioscience, USA) as described previously (Lindholm *et al.*, 2007; Voutilainen *et al.*, 2009). Cells needed to have at least one long neurite and be polygonal to be included in the counting. The results are expressed as mean number of TH-positive cells per section from the 3 sections.

### TH-positive fiber density in the striatum

Analysis of TH-positive fibers in the striatum was performed under blinded conditions. Optical density of TH-positive fibers was measured bilaterally from 3 coronal sections of each rat at approximately the same rostro-caudal sites (AP = +1.2; +0.48 and –0.26 relative to the bregma, according to the rat brain atlas (Paxinos and Watson, 1998). The data are presented as percentage of the lesioned striatum as compared with the intact striatum.

### Experimental Design and Statistical Analysis

Experiments in cultured cells were repeated 3-8 times. The number of animals per group in *in vivo* pERK/pS6 analysis, microdialysis and 6-OHDA model was selected on the basis of the previous experience (Lindholm *et al.*, 2007; Voutilainen *et al.*, 2009) and is indicated in the description of each experiment. The data were subjected to statistical analysis using Student’s t-test or one-way ANOVA with a Dunnett’s or Tukey HSD *post hoc* tests in GraphPad Prism 6 (GraphPad Software Inc., USA) or SPSS^®^ Statistics 22 (IBM SPSS Inc., USA) software. Results were considered statistically significant when P-value was lower than 0.05. Results are presented as Mean±SEM. In the behavioral experiments, TH-immunohistochemical and DAT-immunohistochemical assays data were excluded from the analysis if they exceeded Mean ± 2x Standard Deviation. Vehicle treated rats that were rotating less than 50 net ipsilateral turns in 120 minutes at 2 weeks post lesion were excluded from the results.

## Results

### BT13 stimulates RET phosphorylation and downstream intracellular signaling in immortalized cells

Dopamine neurons are responsive to GDNF (Lin *et al.*, 1993) and NRTN (Akerud *et al.*, 1999) that signal via GFRα1/RET and GFRα2/RET (reviewed in Airaksinen and Saarma, 2002). Therefore, we evaluated the ability of BT13 to stimulate RET phosphorylation and downstream signaling cascades in MG87RET fibroblasts transfected with GFRα1 or GFRα2. We also assessed activation of RET and RET phosphorylation triggered intracellular signaling cascades in response to BT13 in GFP- or expression vector-transfected MG87RET fibroblasts to find out if this compound requires the presence of GFRα1/2 co-receptors to elicit biological effects.

BT13 increased the level of phosphorylated RET (pRET) in GFRα1, GFRα2 and GFP transfected MG87RET fibroblasts in a dose-dependent manner as shown in Figure 1A-C. To quantify the level of pRET in MG87RET/GFRα1, MG87RET/GFRα2 and MG87RET/GFP fibroblasts treated with BT13, GDNF, NRTN and soluble GFRα1 in complex with GDNF, we used RET-ELISA method (Parkash *et al.*, 2008). In line with Western blotting results, BT13 and positive control proteins significantly increased the level of pRET in the cells expressing GFRα1/RET (F (5, 18) = 765.5, P < 0.0001, one-way ANOVA) (Fig 1D), GFRα2/RET (F (5, 18) = 74.30, P < 0.0001, one-way ANOVA) (Fig 1E) and GFP/RET (F (5, 18) = 57.52, P < 0.0001, one-way ANOVA) (Fig. 1F). The effect of BT13 was concentration dependent. Thus, in the cells expressing GFRα1/RET 25 μM BT13 increased the level of pRET by 1.2 fold (P = 0.0116), 50 μM – by 1.4 fold (P < 0.0001) and 100 μM - by 1.9 fold (P < 0.0001). In the cells expressing GFRα2/RET 25 μM BT13 increased the level of pRET by 1.3 fold (P = 0.0088), 50 μM - by 1.6 fold (P < 0.0001) and 100 μM – by 2.1 fold (P < 0.0001). In the cells expressing GFP/RET increased level of pRET was observed in response to 50 μM (1.7 fold increase, P < 0.0001) and 100 μM (2.1 fold increase, P < 0.0001) BT13 (Fig 1 D-F). As expected, GDNF, NRTN and soluble GFRα1/GDNF also increased the level of RET phosphorylation in the cells expressing cognate receptors by 4.6, 2 and 2.6 times, respectively (P < 0.0001, Dunnett’s *post hoc* test for all comparisons). These results were reproduced in at least 4 independent experiments.

**Figure 1:**
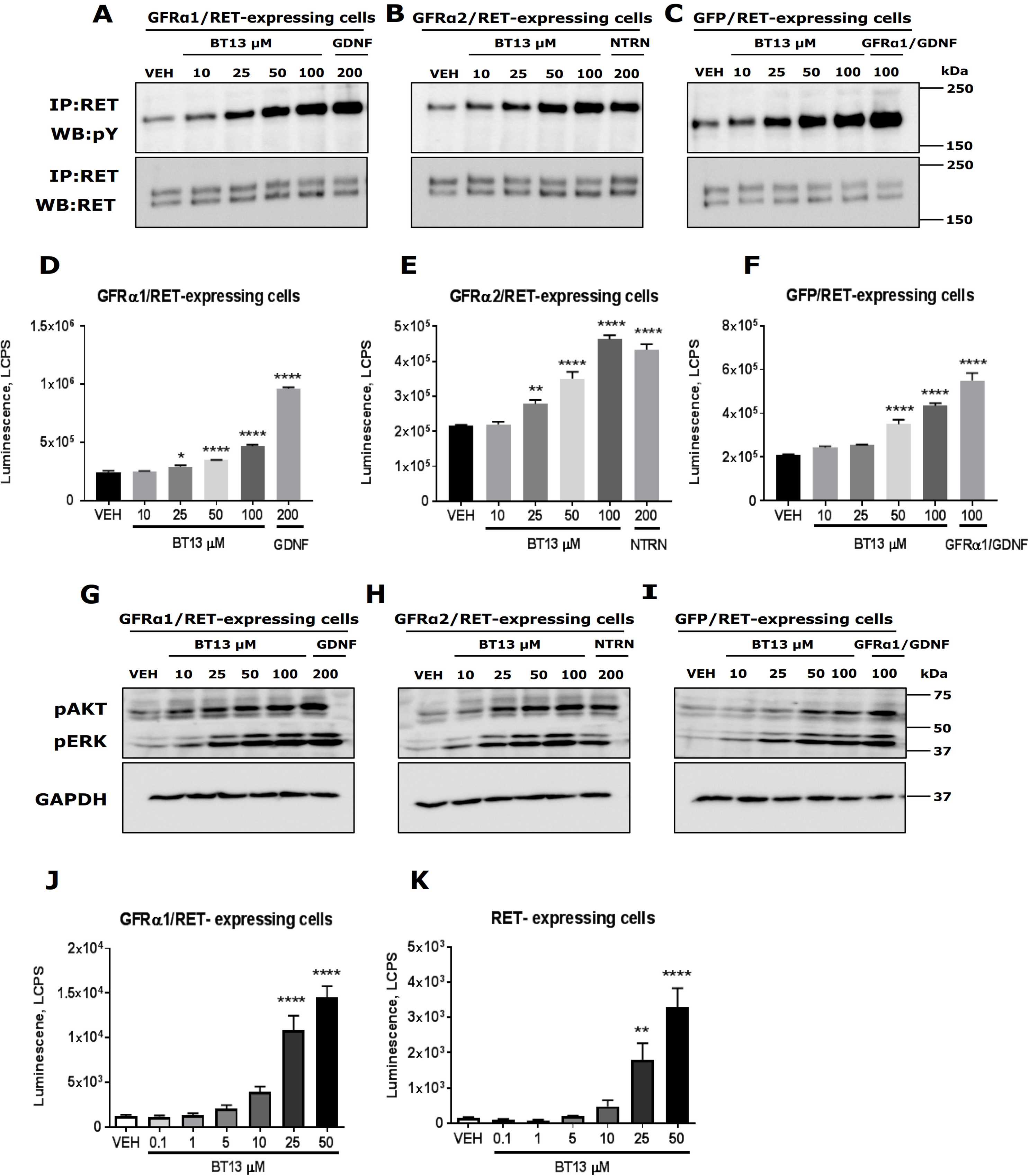
BT13 stimulates phosphorylation of RET (A-C, D-F) and its downstream targets AKT and ERK (D-F, G-I) in immortalized MG87RET fibroblasts. Assessment of RET (A-C) and AKT/ERK phosphorylation by Western blotting (G-I), phospho-RET-ELISA assay to quantify the level of RET phosphorylation (D-F) and activation of luciferase reporter controlled by RET via ERK (J,K). Effect of BT13 in MG87RET fibroblasts transfected with GFRα1 (A, D, G, J), GFRα2 (B, E, H) and GFP (C, F, I, K). Concentrations of GDNF, NTRN and soluble GFRa1/GDNF complex used as positive controls for GFRα1, GFRα2 and GFP transfected MG87RET cells, respectively, are provided in ng/ml. VEH - vehicle, IP - immunoprecipitation, WB - Western blotting, GAPDH – glyceraldehyde-3-phosphate dehydrogenase, a house-keeping protein, loading control. * P < 0.05, ** P < 0.01, **** P < 0.0001, one-way ANOVA with Dunnett’s *post hoc* test, N=3-4.

GDNF- and NRTN-dependent autophosphorylation of RET causes activation of downstream signaling pathways. Since GDNF-dependent AKT phosphorylation is crucial for neuronal survival and ERK phosphorylation for neurite outgrowth, we assessed the phosphorylation of RET downstream targets AKT and ERK in GFRα1/RET, GFRα2/RET and GFP/RET-expressing cells. BT13 clearly activated both downstream targets in presence (GFRα1- and GFRα2-transfected) and absence (GFP-transfected) of GFRα co-receptors (Fig 1G-I). Cells responded in the expected manner to GDNF and NTRN used as positive controls. To provide a quantitative estimate on the level of intracellular signaling cascade stimulation in response to BT13, we used luciferase reporter gene-based assay reflecting the level of ERK activation (Sidorova *et al.*, 2010). In line with Western blotting data (Fig 1G-I), RET phosphorylation data (Fig 1A-C, D-F) and published reports (Sidorova *et al.*, 2017), BT13 increased luciferase activity in a concentration dependent manner in reporter cell lines expressing GFRα1/RET (F (6, 14) = 43.78, P < 0.0001, one-way ANOVA) and RET (F (6, 14) = 19.85, P < 0.0001, one-way ANOVA). BT13 in concentrations 25 and 50 μM increased luciferase activity in GFRα1/RET expressing cells by 8.6 fold and 11.5 fold, respectively (P < 0.0001), and in RET expressing cells by 11.9 fold (P =0.0040) and 21.7 fold, respectively (p<0.0001, *post hoc* comparisons with Dunnett’s test in all cases) (Fig. 1J-K). In this assay GDNF usually increases luciferase activity in GFRα1/RET expressing cells by 50-100 fold (Sidorova *et al.*, 2010, 2017; Saarenpää *et al.*, 2017).

### BT13 supports the survival of cultured dopamine neurons

GDNF is a well-known neurotrophic factor supporting the survival and maintenance of embryonic dopamine neurons in culture (Lin *et al.*, 1993). Therefore, we assessed whether BT13 also promotes the survival of cultured dopamine neurons. BT13 and GDNF increased the number of TH-positive neurons in cultures (RM ANOVA F (2.35, 16.46) = 6.55, P = 0.0062). On 5^th^ DIV the number of TH-positive cells in the wells treated with 0.1 μM and 1 μM BT13 increased by 1.4 fold (P < 0.0001) and 1.5 fold, respectively (P = 0.0002, Dunnett’s *post hoc* test in both cases), compared to vehicle-treated wells. GDNF (10 ng/ml) increased the number of TH-positive cells in culture by 1.8 fold (P = 0.0207, Dunnett’s *post hoc* test) (Figure 2A, B). Important to note, that the bell-shaped dose-response curves are common for neurotrophic factors in general and for GDNF in particular (Hou *et al.*, 1996; Saarenpää *et al.*, 2017). Therefore, the lack of increase in dopamine neuron survival in response to the higher doses of BT13 is expected.

**Figure 2:**
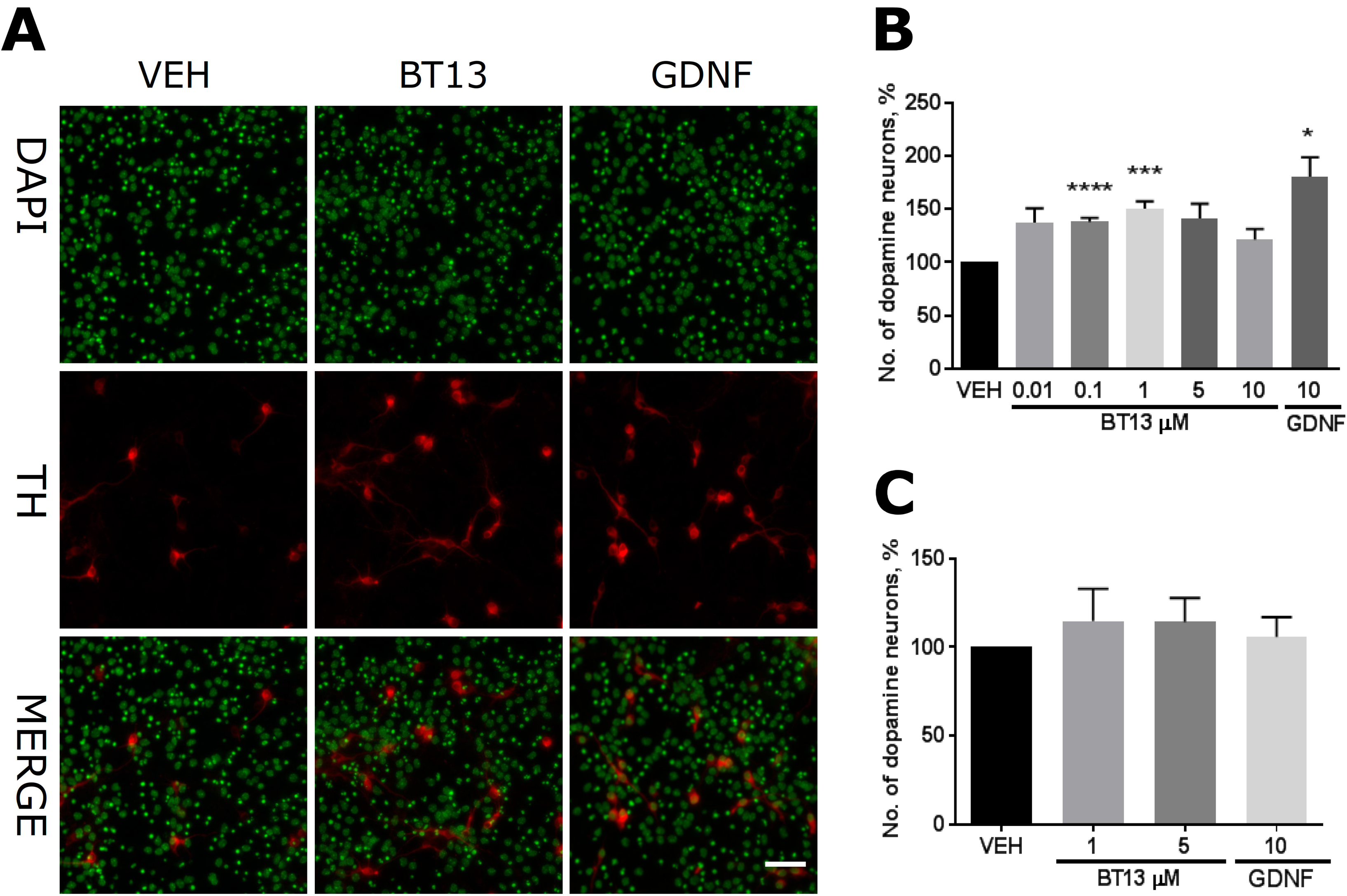
BT13 similarly to GDNF promotes the survival of cultured primary midbrain dopamine neurons from wild-type (A, B), but not from RET knock-out mice (C). (A) Representative images of mouse E13.5 wild-type midbrain cultures treated with vehicle, BT13 and GDNF probed with anti-TH antibody, pseudocolors. (B) The number of TH-positive cells in the wild-type midbrain cultures on 5^th^ DIV normalized to the total number of cells in the culture and presented as percentage of vehicle treated samples. (C) The number of TH-positive cells in the RET knock-out midbrain cultures on 5^th^ DIV normalized to the total number of cells in the culture and presented as percentage of vehicle treated samples, average from 3 experiments. Concentration of GDNF used as a positive control is provided in ng/ml. VEH - Vehicle. * P < 0.05, *** P < 0.001, **** P < 0.0001, RM ANOVA with Dunnett’s *posthoc* test, N = 8. Scale bar – 50 μm.

To ensure that the effect of BT13 on the survival of dopamine neurons is RET dependent, we assessed the survival of RET knock-out embryonic midbrain neurons treated with BT13 and GDNF. The number of TH-positive cells in the cultures of RET knock-out midbrains on 5 ^th^ DIV remained unchanged in response to either BT13 or GDNF (Fig 2C).

We also evaluated the neuroprotective properties of BT13 in dopamine neurons in *in vitro* model of Parkinson’s disease in duplicates in a single experiment. To induce the death of dopamine neurons, rat E15 embryonic midbrain cultures on 6^th^ DIV were treated with MPP^+^ and BT13. On 8^th^ DIV the number of remaining TH-positive neurons was assessed. In these conditions BT13 in concentrations 2, 20 and 60 nM increased the number of remaining TH-positive neurons by 15%, 16% and 13% in comparison to vehicle, respectively. In the same experiment BDNF increased the number of remaining TH-positive neurons after MPP^+^ treatment by 22%.

### BT13 stimulates intracellular signaling important for neuronal survival and regeneration in the mouse striatum

As BT13 activated ERK and AKT intracellular signaling cascades in immortalized cells, we assessed if it can do so also *in vivo* by measuring the levels of phosphorylated ERK and ribosomal protein S6 (downstream target of AKT) in the mouse striatum. Each mouse was injected with vehicle into the right striatum and BT13 or GDNF into the left striatum (Fig 3A). The level of pERK and pS6 staining in vehicle-treated striata was the same in all treatment groups (P > 0.05, one-way ANOVA). BT13 at the dose of 750 μg significantly increased phosphorylation of ERK and S6 as compared to the vehicle treated striatum (t(2) = 4.38, P = 0.048 and t(3) = 3.47, P = 0.040, respectively, paired two-tailed Student’s t-test) (Fig 3 B, C). GDNF at the dose of 5 μg elevated levels of both pERK (t(2) = 6.003, P = 0.0266) and pS6 (t(2) = 15.23, P = 0.0043) and at the dose of 10 μg increased phosphorylation of ERK (t(3) = 6.194, P = 0.0085) in the mouse striata.

**Figure 3:**
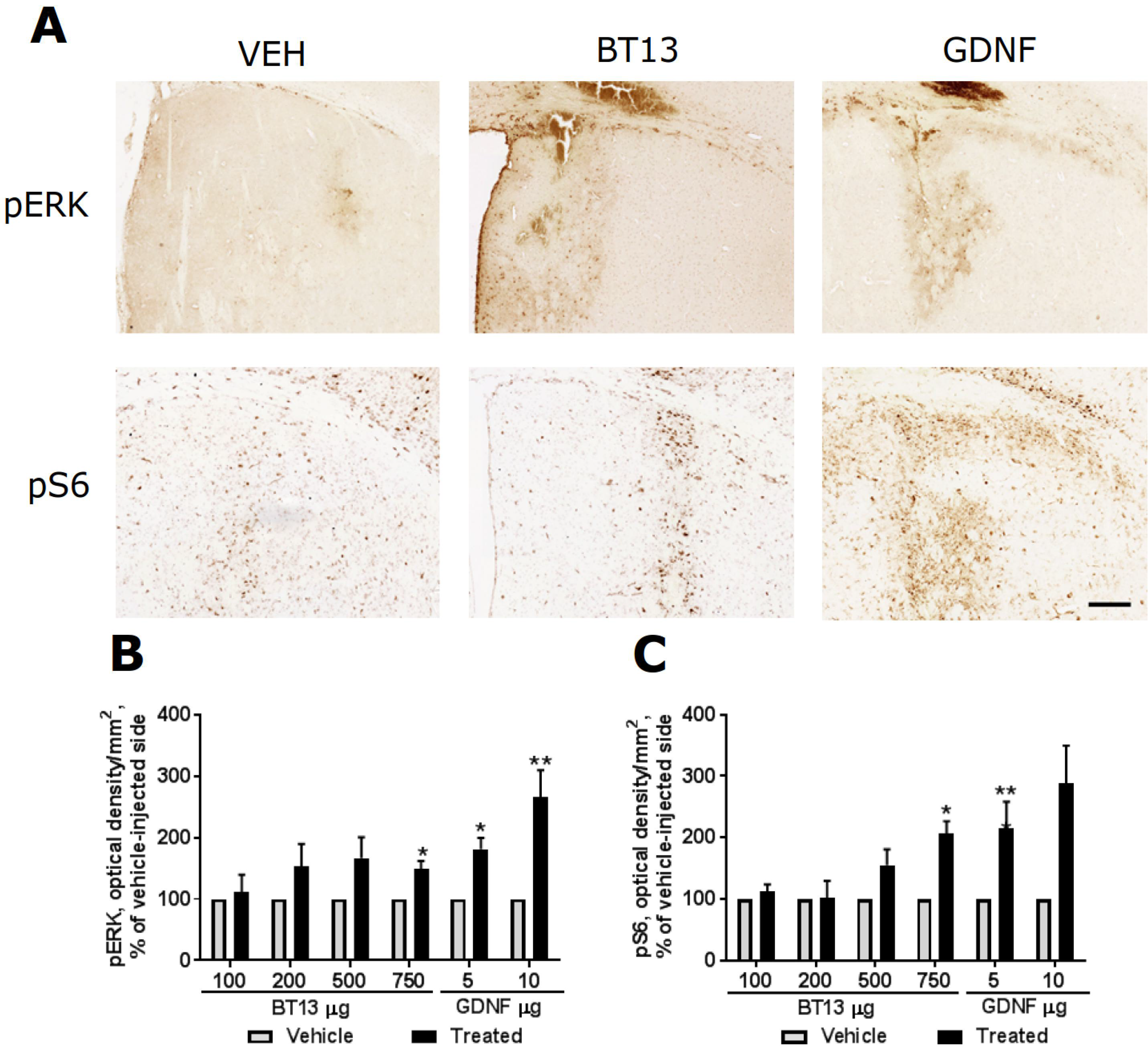
BT13 activates intracellular signaling cascades responsible for survival and regeneration of neurons *in vivo*. (A) Representative images of coronal sections from mouse brains probed with anti-phospho-S6 (pS6, lower pictures) and anti-phospho-ERK (pERK, upper pictures) antibodies. GDNF dose - 10 μg, BT13 dose - 750 μg. VEH – vehicle. (B) Relative optical density of pERK immunostaining in the dorsal striatum in different treatment groups presented as percent of vehicle treated side. (C) Relative optical density of pS6 immunostaining in the dorsal striatum in different treatment groups presented as percent of vehicle treated side. Statistical significance for the differences between vehicle and BT13/GDNF-injected sides was determined by paired two-tailed Student’s t-test (* P < 0.05; ** P < 0.01), N = 3-4/group. Scale bar: 200 μM

### BT13 stimulates release of dopamine in the mouse striatum

We further tested the ability of BT13 to induce dopamine release in the mouse striatum using brain microdialysis, as GDNF has been shown to increase striatal dopamine release (Hebert *et al.*, 1996; Pothos *et al.*, 1998; Grondin *et al.*, 2003; Gomes *et al.*, 2006). BT13 treatment was initiated 30 minutes after the beginning of the experiment (*i.e.* after obtaining 4 stable baseline values) and we estimate that it took approximately 60 minutes for the solution to reach the striatum via tubing. The infusion of 50 μM BT13 resulted in a clear increase (F (17, 72) = 5.30, P < 0.0001, one-way ANOVA) in extracellular dopamine level in the striatum (Fig. 4) that was observed during 105-165 minutes (30 min vs 105 min P=0.0160; 30 min vs 120 min P=0.0011; 30 min vs 135 min P=0.0013; 30 min vs 150 min P=0.0015; 30 min vs 165 min P=0.0091, one-way ANOVA with Dunnett’s *post hoc* test) from the beginning of the experiment (or approximately 75-135 minutes after the beginning of BT13 infusion). The effect reached the maximum at 90 minutes from the initiation of BT13 infusion and remained at practically the same level for the next 30 minutes, followed by a gradual decrease. In vehicle-treated mice the level of extracellular dopamine remained stable during the whole experiment (data not shown).

**Figure 4:**
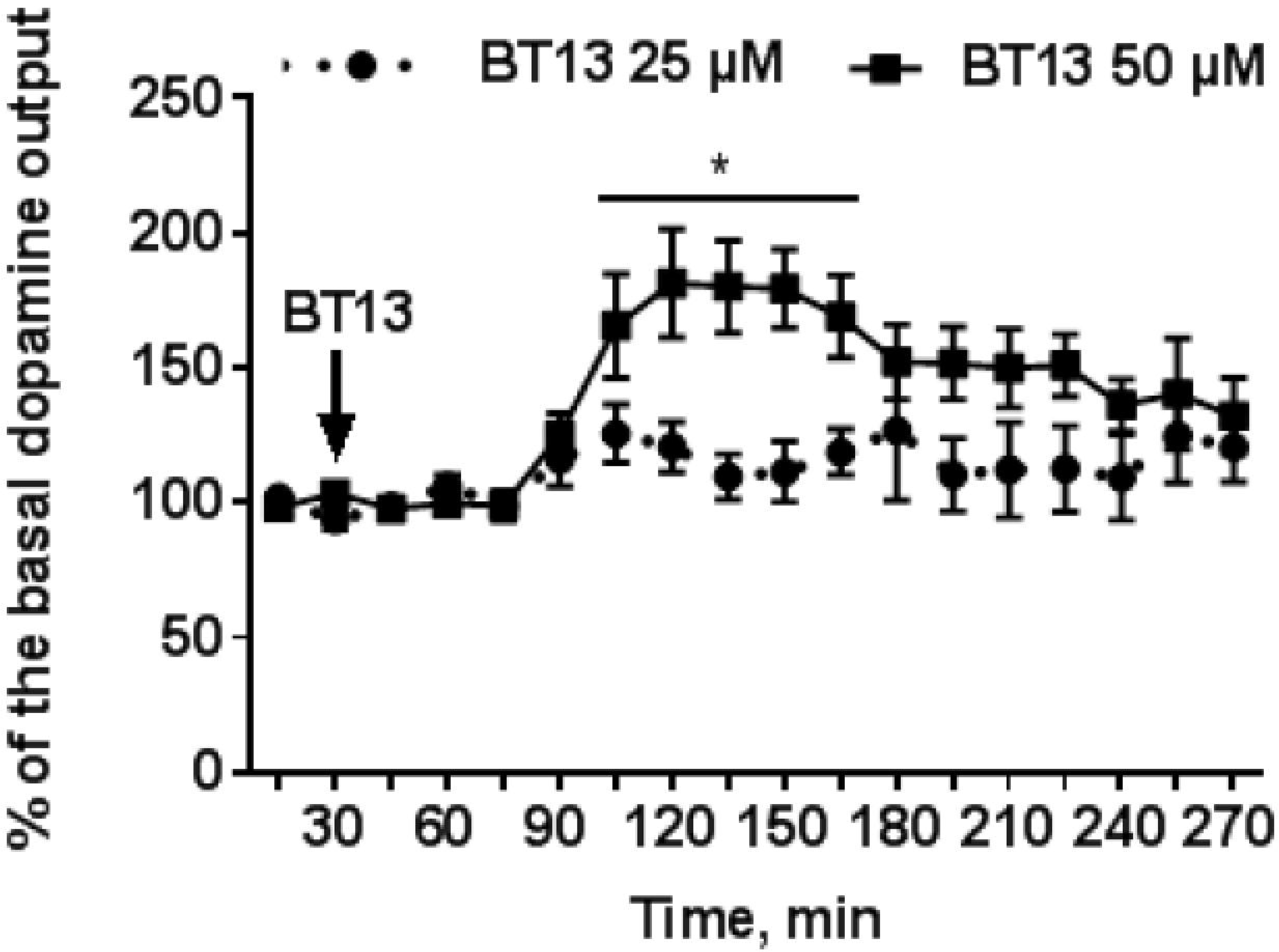
Effect of continuous infusion of BT13 on extracellular dopamine level in the mouse dorsal striatum. BT13 in 50 μM concentration increased extracellular dopamine level, while the lower concentration (25 μM) produced a small and statistically insignificant effect. * P < 0.05, one-way ANOVA with Dunnett’s *post-hoc* test, Mean ± SEM, N = 5/group

### BT13 alleviates motor deficits in 6-OHDA rat model of Parkinson’s disease

The neuroprotective effect of BT13 was compared with that of GDNF in a 6-OHDA-model of Parkinson’s disease in rats. The scheme of the experiment is presented in Fig. 5A. In vehicle treated rats, amphetamine induced a strong ipsilateral turning behavior at all tested time points indicating unilateral damage to dopaminergic system on the 6-OHDA lesioned side of the brain (Fig. 5B-D). The number of net ipsilateral turns was similar in all treatment groups at 2 weeks post lesion (Fig 5B). At 4 weeks post lesion, the number of net ipsilateral turns in animals treated with BT13 or GDNF was reduced significantly in comparison to the vehicle (F(2.53) = 7.344; P = 0.002; one-way ANOVA) (Fig. 5C). Tukey HSD *post hoc* test revealed that the number of turns was significantly lower in BT13 3-6 μg/24 h (136.3 ± 30.1; P = 0.013) and GDNF 3 μg/24 h (46.9 ± 23.3; P = 0.001) groups as compared to the vehicle group (350.8 ± 106.1). The effect of BT13 3-6 μg/24 h was significant and comparable with the effect of GDNF 3 μg/24 h also 6 weeks after the lesion (F(2.51) = 5.882; P=0.005; one-way ANOVA) (Fig. 5D). According to Tukey HSD *post hoc* test the number of turns was significantly lower in BT13 3-6 μg/24 h (13.1 ± 10.1; P = 0.011) and GDNF 3 μg/24 h (−11.8 ± 8.1; P = 0.007) groups as compared to the vehicle group (171.1 ± 93.1).

**Figure 5:**
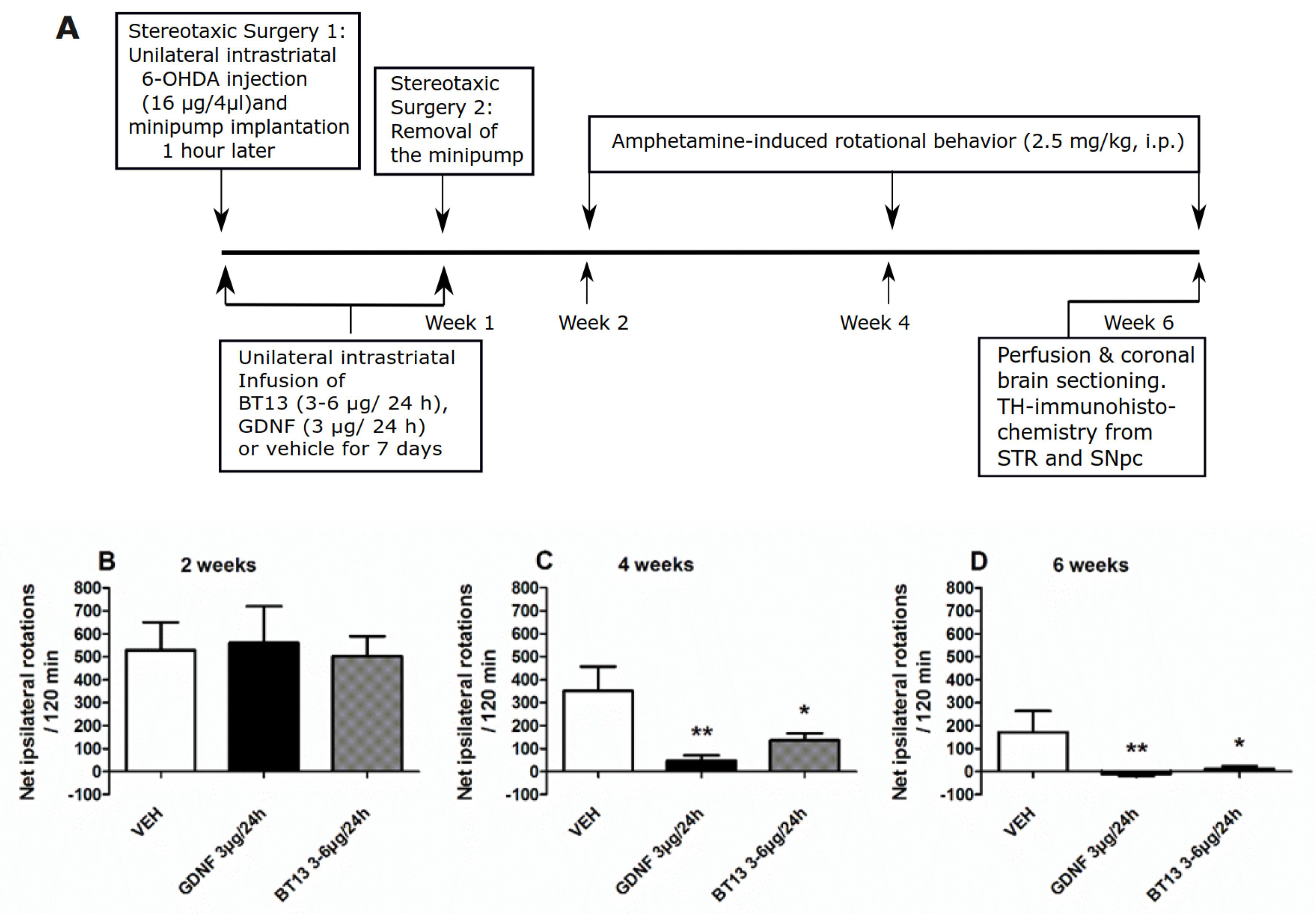
Amphetamine-induced turning behavior is normalized by BT13 and GDNF in a time-dependent manner. (A) Scheme of the *in vivo* neuroprotection experiment. (B) Effect of BT13 and GDNF infusions on amphetamine-induced turning behavior at 2 weeks, (C) 4 weeks, and 6 weeks post lesion. VEH – vehicle. * P < 0.05, ** P < 0.01 as compared to the vehicle, one-way ANOVA with Tukey HSD *post-hoc* test, Mean ± SEM, vehicle N = 11-12, GDNF 3μg/24h N = 15-16, BT13 3-6μg/24h N = 25-27.

Noteworthily, in line with previously published data on this particular toxin administration scheme (Penttinen *et al.*, 2016), we observed spontaneous recovery in the turning behavior in vehicle-treated rats at 4 and 6 weeks post lesion (350.8 ± 106.1 and 171.1 ± 93.1 net ipsilateral turns per 120 minutes, respectively, vs. 529.8 ± 119.8 net ipsilateral turns per 120 minutes at 2 weeks post lesion; Student’s paired two-tailed t-test for 2 weeks vs. 4 weeks post lesion t(11) = 1.895, P = 0.085; for 2 weeks vs. 6 weeks post lesion t(10) = 2.460, P = 0.034; for 4 weeks vs. 6 weeks post lesion t(10) = 2.756, P = 0.020).

### Effect of BT13 on TH-immunoreactive neurons in the SNpc and the density of TH-immunoreactive fibers in the striatum of 6-OHDA lesioned rats

To assess the extent of the lesion in the neuroprotection study, morphological analysis of nigral and striatal sections stained with TH antibody was performed after the last behavioral test. The single unilateral injection of 6-OHDA into the dorsal striatum resulted in a relatively mild degeneration of dopamine neurons: in vehicle treated rats, the density of TH-positive fibers in the striatum was reduced by 55.5 ± 3.9% in comparison to the intact side (Fig. 6A, D). As anticipated, GDNF infusion significantly protected TH-positive fibers in the lesioned striatum (43.1 ± 3.1% loss in TH-immunoreactivity) as compared to vehicle treatment (P = 0.024; Tukey HSD *post-hoc* test after one-way ANOVA F(2,51) = 3.708; P = 0.032) (Fig. 6D). BT13 also seemed to have an effect in protecting TH-positive fibers (47.9 ± 2.1% loss in TH-immunoreactivity) but this effect did not reach statistical significance when compared to the vehicle (P = 0.180; Tukey HSD *post hoc* test). The number of TH-positive cell bodies in the SNpc on 6-OHDA lesioned side was reduced by 27.2 ± 10% in vehicle treated rats (Fig. 6B, E). Intrastriatal infusion of BT13 or GDNF was unable to protect TH-positive cells in the SNpc (Fig. 6E). Since GDNF overexpression or injection into striatum can downregulate TH-expression (Rosenblad *et al.*, 2003; Georgievska *et al.*, 2004; Salvatore *et al.*, 2004), we analysed the density of DAT-immunoreactive fibers in the striatum of GDNF and BT13 treated rats (Fig. 6C, F). The density of DAT-positive fibers was comparable to the density of TH-positive fibers in all treatment groups suggesting unaltered TH expression in this study protocol (Fig. 6F).

**Figure 6:**
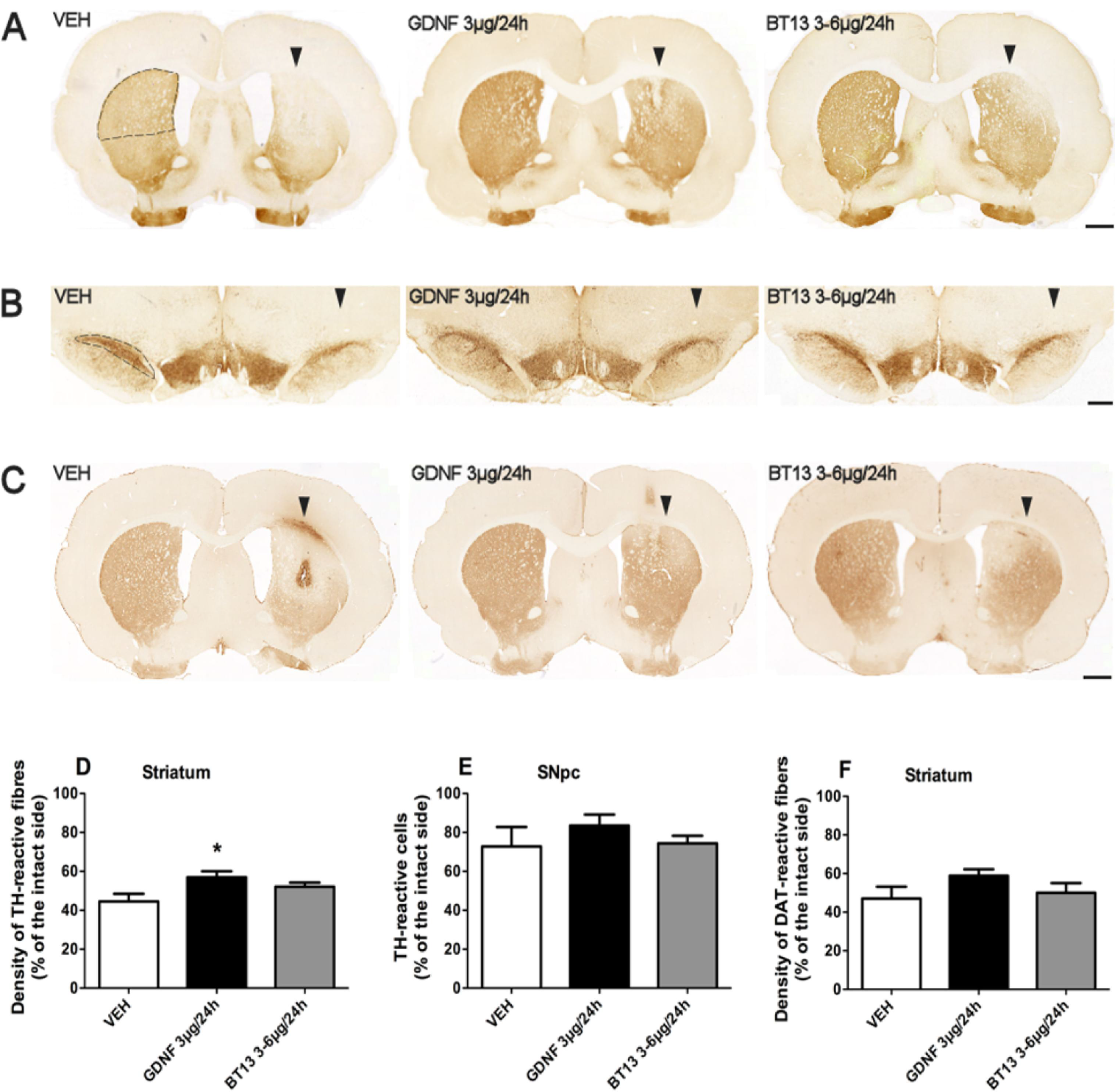
Effect of BT13 and GDNF on the density of TH and DAT-immunoreactive fibers in the dorsal striatum and the number of TH-immunoreactive cells in the SNpc in 6-OHDA lesioned rats. (A) Representative pictures of TH immunohistochemical staining in the striatum, (B) TH-immunoreactive cells in the central SNpc, and (C) DAT immunoreactivity in the striatum in different treatment groups 6 weeks post lesion. (D) Quantification of the density of TH-immunoreactive fibers in the dorsal striatum. (E) Quantification of the number of TH-immunoreactive cells in the SNpc. (F) Quantification of the density of DAT-immunoreactive fibers in the dorsal striatum. The dashed line in (A) shows the site of optical density measurements in the dorsal striatum and in (B) the site of stereological cell counting. Scale bar in (A) and (C) is 1 mm and in (B) 0.5 mm. The lesion-side is denoted with arrowheads. VEH – vehicle. * P < 0.05 as compared to the vehicle, one-way ANOVA with Tukey HSD *post hoc* test, Mean ± SEM. In TH staining (D, E) vehicle N = 11-12, GDNF 3μg/24h N = 15-16, BT13 3-6μg/24h N = 24-25; in DAT staining (F) vehicle N = 8, GDNF 3μg/24h N = 13, BT13 3-6μg/24h N = 19.

## Discussion

In the present study, we demonstrated that a small molecule RET agonist, BT13, promotes the survival of embryonic dopamine neurons *in vitro* and protects them from MPP^+^ induced toxicity. Short-term administration of BT13 into the striatum of intact mice activated intracellular signaling cascades important for the survival and regeneration of dopamine neurons and increased extracellular dopamine levels. In unilateral 6-OHDA rat model of Parkinson’s disease, BT13 improved functional recovery of neural circuits controlling movements and reduced motor imbalance caused by toxin-induced degeneration of dopamine neurons. This effect was accompanied by a trend to increase in the density of TH-immunoreactive dopaminergic fibers in the striatum of 6-OHDA lesioned rats in response to BT13.

In immortalized cells BT13 had similar efficacy to GFLs in RET, ERK and AKT phosphorylation assay (Fig 1 A-F), however, in integral luciferase assay the effect of BT13 was much more modest in comparison to GDNF or soluble GFRα/GDNF complex (Fig 1 G-I, (Sidorova *et al.*, 2010, 2017). This can be at least in part explained by quick metabolism of BT13 (Sidorova *et al.*, 2017). At the same time, the efficacy of BT13 (100 nM and 1 μM) was comparable with that of GDNF in its ability to support the survival of cultured embryonic dopamine neurons (Fig. 2), to attenuate amphetamine-induced turning behavior in 6-OHDA model of Parkinson’s disease in rats (at approximately 0.5-1 mM concentration,Fig. 5), and to increase phosphorylation of ERK and ribosomal protein S6 (Fig. 3) in the striatum of naïve mice at the dose of 750 μg. It is important to note, that the concentrations of the positive control proteins in all experiments were selected on the basis of previous results and biological activity in screening assay in order to produce clear effects in studied systems (Trupp *et al.*, 1996; Sidorova *et al.*, 2010, 2017; Runeberg-Roos *et al.*, 2016; Saarenpää *et al.*, 2017). Nevertheless, the potency of BT13 was two-four orders of magnitude lower: GDNF supported the survival of cultured dopamine neurons in approximately 0.3 nM concentration and reduced the number of amphetamine-induced turns at about 8 μM concentration.

Importantly, the survival of the primary dopamine neurons was tested in much lower GDNF and BT13 concentrations than the biochemical effects in immortalized cells. Artificial cell systems differ from primary neurons in many ways. In particular, the stoichiometry of relevant receptors, intracellular mediators and other factors may differ and hereby regulate sensitivity to agonists. In addition, limited aqueous solubility of BT13 made it complicated to test a higher concentration (above 100 μM) of the compound in murine fibroblasts. Perhaps in immortalized cells the full potential of BT13 to promote phosphorylation of RET and intracellular signaling cascades has not been achieved because of its low solubility in the cell culture medium. In addition to activation of RET and downstream signaling pathways in cell lines, injection of BT13 into the intact mouse striatum, similarly to GDNF, activated intracellular ERK and AKT signaling cascades (Fig. 3). Thus, BT13 appears to be able to activate the same downstream signaling pathways in the striatum as GDNF.

Microdialysis data show that infusion of BT13 into the striatum increases extracellular dopamine level (Fig. 4) further indicating its ability to activate relevant signaling pathways in the brain. Notably, in transgenic mice overexpressing GDNF from the native locus, enhanced dopamine release in the striatum is detected (Kumar *et al.*, 2015). It is important to note that the ability of BT13 to elevate extracellular dopamine level may not correlate with its neurotrophic activity. The effect on dopamine level is possibly mediated through the inhibition of A-type potassium channels that GDNF is known to regulate (Yang *et al.*, 2001), but it may also be the consequence of DAT regulation that was recently characterized for GDNF (Zhu *et al.*, 2015; Kopra *et al.*, 2017). The fact that the extracellular dopamine level started to decrease after some time even though the infusion continued suggests activation of homeostatic mechanisms. These mechanisms might include alterations in the availability or activity of GDNF receptors and/or DAT (Eriksen *et al.*, 2010).

In 6-OHDA rat model of Parkinson’s disease, both BT13 and GDNF improved functional motor recovery indicated by reduced amphetamine-induced turning behavior (Fig. 5B-D). This may be due to prevention of toxin entry into dopamine neurons via influencing DAT activity by BT13, due to dopamine mobilizing effect of BT13 as seen in the microdialysis experiment, or due to its neuroprotective effect on lesioned dopamine neurons. The first mentioned two options are, however, unlikely because BT13 was shown to mobilize dopamine after an acute administration whereas the amphetamine-induced turning behavior was reduced only at 4 weeks post lesion (i.e. 3 weeks after the cessation of the BT13 infusion). We also observed no changes in DAT activity in response to BT13 in an *in vitro* assay (Sidorova *et al.*, 2017). Thus, direct inhibition of 6-OHDA entry into neurons is improbable, although indirect interactions cannot be completely excluded on the basis of these results.

GDNF is known to protect and restore nigrostriatal dopamine neurons against toxin-induced damage in animal models of Parkinson’s disease and preserve normal motor functions in 6-OHDA lesioned animals (Hoffer et a., 1994; Kirik et al., 2000). In the present study, GDNF was able to protect TH-positive dopaminergic innervation in the striatum against 6-OHDA induced degeneration (Fig. 6D). There was also a trend for neuroprotection of TH-immunoreactive fibers in the striatum after BT13 administration, but this effect remained insignificant. DAT-immunostaining showed comparable results for dopaminergic degeneration at the level of the striatum supporting the results from the TH-immunohistochemical analysis (Fig. 6F). We did not see statistically significant effects on the number of TH-positive neurons in the SNpc in response to GDNF or BT13 (Fig. 6E). It should be noted that the concentration of dopamine in striatum may not correlate well with the number of cell bodies in SNpc. Indeed, in patients with Parkinson’s disease at the moment of diagnosis, the degeneration of axons and the drop of dopamine levels in the striatum are more pronounced than the reduction in the number of dopamine neurons cell bodies (Cheng *et al.*, 2010). It can also be speculated that the doses of BT13 and GDNF (or the biological activity of the particular batch of GDNF) were too low to be able to protect dopaminergic cell bodies in the SNpc after intrastriatal infusion. Furthermore, Marco et al. (Marco *et al.*, 2002) reported downregulation of RET and GFRα1 receptors during the first week after intrastriatal 6-OHDA lesion in rats. As the neurotrophic treatments in our study occurred during this period of time when the receptors were downregulated, we were not able to detect as prominent neuroprotective effect of BT13 and GNDF at the level of TH and DAT immunostainings as expected. In contrast to GDNF, BT13 spreads well in tissues and is able to penetrate through tissue barriers (Sidorova *et al.*, 2017). Therefore, we cannot exclude the effects of BT13 on the expression of TH and DAT on the contralateral hemisphere. We express the density of TH- and DAT-immunoreactive fibers in percentage of the contralateral side. Thus, RET activation on the contralateral side resulting in enhanced axonal sprouting could have masked the neuroprotective effect of BT13 on dopamine neurons *in vivo*. Nevertheless, the infused concentration of BT13 was relatively low, and it is unclear if a biologically effective amount of BT13 was able to reach the contralateral side of the brain.

In general, the model we used in this study was characterized by a relatively mild lesion of dopamine neurons and significant spontaneous functional recovery. Although in this partial lesion model BT13 did not show significant neuroprotection at the level of TH immunohistochemistry, the significant functional improvement in the behavioral tests (Fig 5B-D) together with activation of similar intracellular signaling cascades (Fig. 3) suggests similar neuroprotective mechanism of action for BT13 and GDNF *in vivo.*

Noteworthily, the pharmacological properties of BT13 still require improvement. Although BT13 is able to cross the BBB (on average 55% of the compound reach the brain (Sidorova *et al.*, 2017)), its aqueous solubility is relatively low (100 μM). We used propylene glycol to dissolve BT13, however, after infusion of the compound into aqueous extracellular environment its low solubility could lead to partial precipitation and reduction of its free concentration in the brain. In addition, BT13 is metabolically unstable (Sidorova et al., 2017), thus the dose, administration route and delivery scheme for this compound may require careful considerations in the future.

Taken together, our data offer a proof of principle for small molecule synthetic RET agonists in animal model of Parkinson’s disease. Our first generation compound BT13 has promising effects on dopaminergic system both *in vitro* and *in vivo*, but substantial optimization is needed to promote it to clinical use. We are currently improving the potency and pharmacological properties of BT13 using methods of medicinal chemistry to develop a lead compound for clinical trials in patients with Parkinson’s disease.

## Acknowledgements

We thank Solvay Pharmaceuticals BV for planning and financing the experiments with 1-methyl-4-phenylpyridinium (MPP ^+^) challenged dopamine neurons and Neuron Experts company (Marseilles, France) for performing these experiments. We acknowledge the help of Dr. Satu Kuure with experiments with RET knock-out dopamine neurons. We thank Dr. Harri Jäälinoja from Light Microscopy Unit of University of Helsinki for guidance in establishing of automated quantification protocol for analysis of the survival of dopamine neurons. The Nuclear Magnetic Resonance (NMR) facility at the Institute of Biotechnology, University of Helsinki supported by Biocenter Finland and Helsinki Institute of Life Science (HiLIFE) is acknowledged for verification of BT13 structure. We are grateful to Kati Rautio, Marjo Vaha, Laura Salminen, Sascha Gromnitza, Elisa Piranen, Jenni Montonen, Leo Jakman, Jette Sakki, Charlotte Zeitler, and Laura Bravo-Burguillos, for their excellent assistance with the experimental procedures.

## Author contributions

AKM, J-MR, JK, TV, IK, NP, YS collected, analyzed and interpreted the data; ER, TPP, MV, RKT, MBB, MK, YS, MS designed the study, analyzed and interpreted the data; AKM, J-MR, JK, TV, ER, TPP, MV, RKT, YS, MS drafted the manuscript. All authors approved the final version of the manuscript.

## The paper explained

### Problem

Parkinson’s disease (PD) is the second most common neurodegenerative disorder. Diagnostic motor symptoms of PD are caused by the degeneration and death of dopamine neurons in the brain. Existing treatments alleviate PD symptoms, but fail to support injured dopamine neurons or by other words the curative therapy for PD does not exist.

### Results

In the present study we show that BT13, the small molecule agonist of receptor tyrosine kinase RET, activates intracellular events important for the well-being of dopamine neurons in *in vitro* and *in vivo* settings, promotes the survival of dopamine neurons cultured in the dish, stimulates the function of dopamine system and alleviates motor manifestations of PD in experimental animals.

### Impact

Our data indicate that BT13 can pave a way to the future development of curative therapy against PD.

## Expanded View section

### Detailed description of materials and methods

#### Cell Lines

MG87 RET murine fibroblasts stably transfected with RET oncogene (Eketjäll *et al.*, 1999). MG87RET fibroblasts stably transfected with GFRa1-expressing plasmid or empty vector and a luciferase reporter gene system (PathDetect detect Elk-1 trans-Reporting system, Stratagene/Agilent Technologies, USA) to detect activation of mitogen-activated protein kinase (MAPK) signaling pathway (Sidorova *et al.*, 2010).

#### Proteins

Human recombinant GDNF (hGDNF) for *in vitro* experiments was produced in mammalian CHO cells and obtained from Icosagen (Estonia), hGDNF for *in vivo* studies was produced in *E.coli* and was purchased from PeproTech (USA).

#### BT13

The synthetic route for the compound BT13 is given below

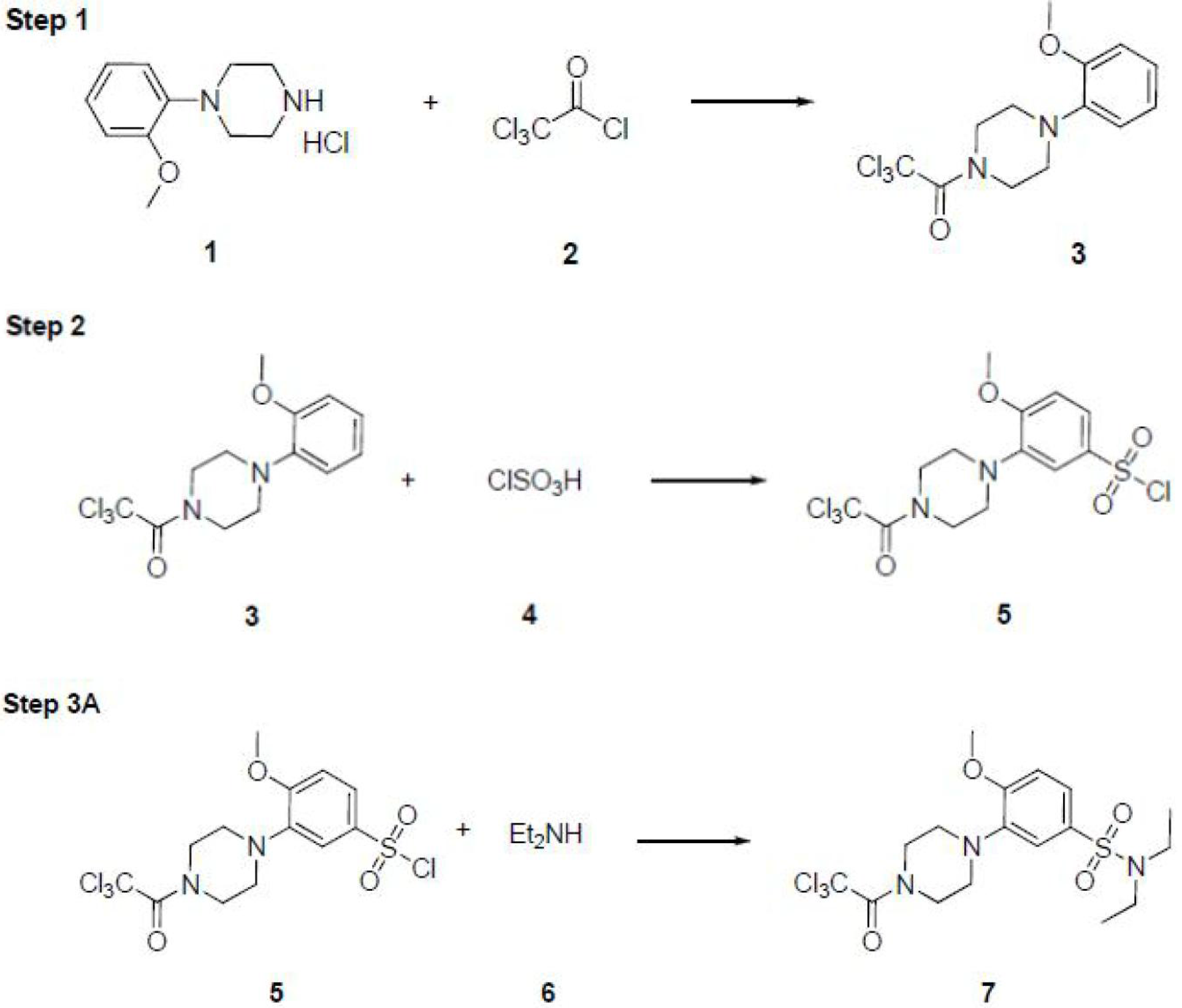

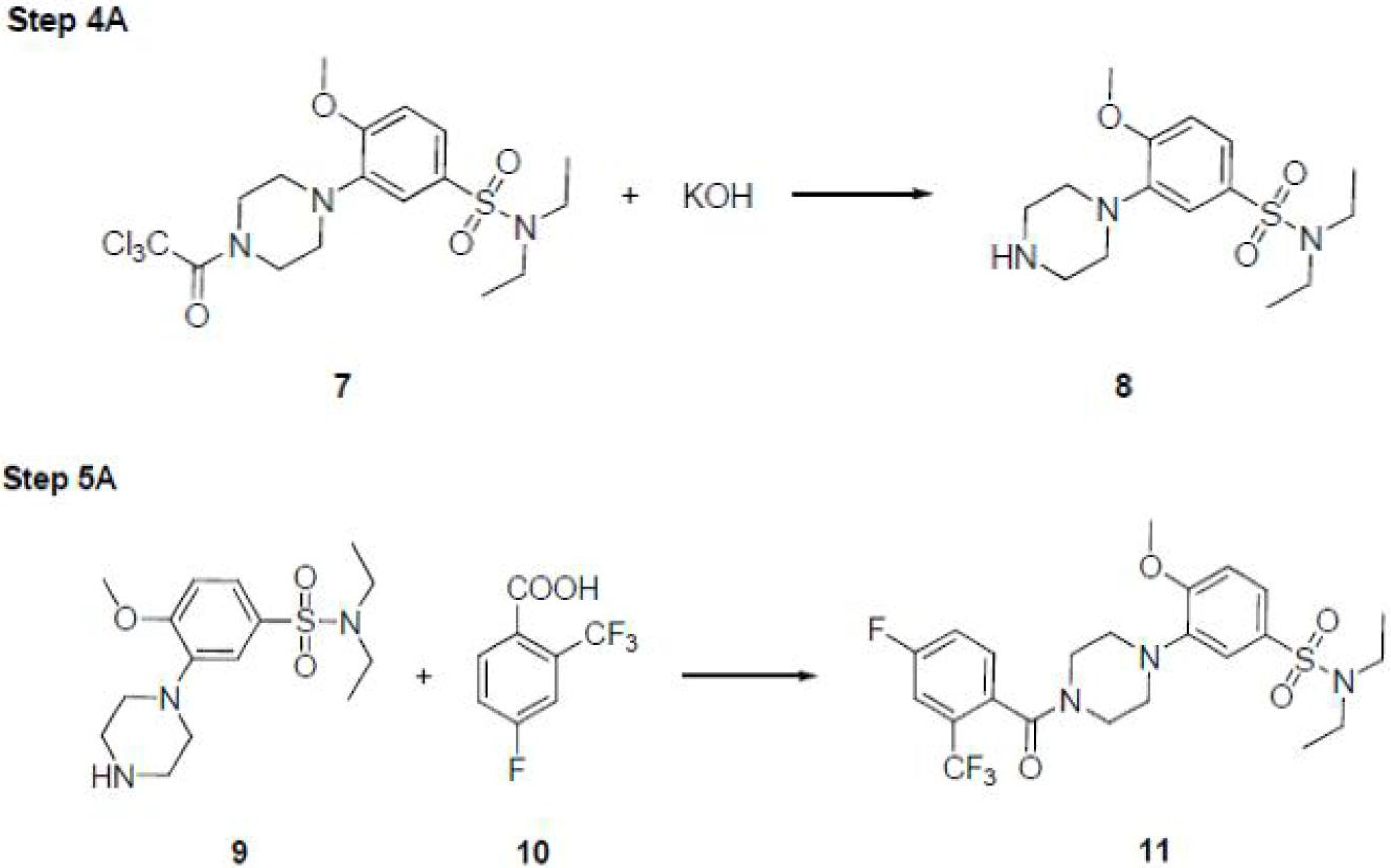

The individual steps of the synthesis were carried out as follows.

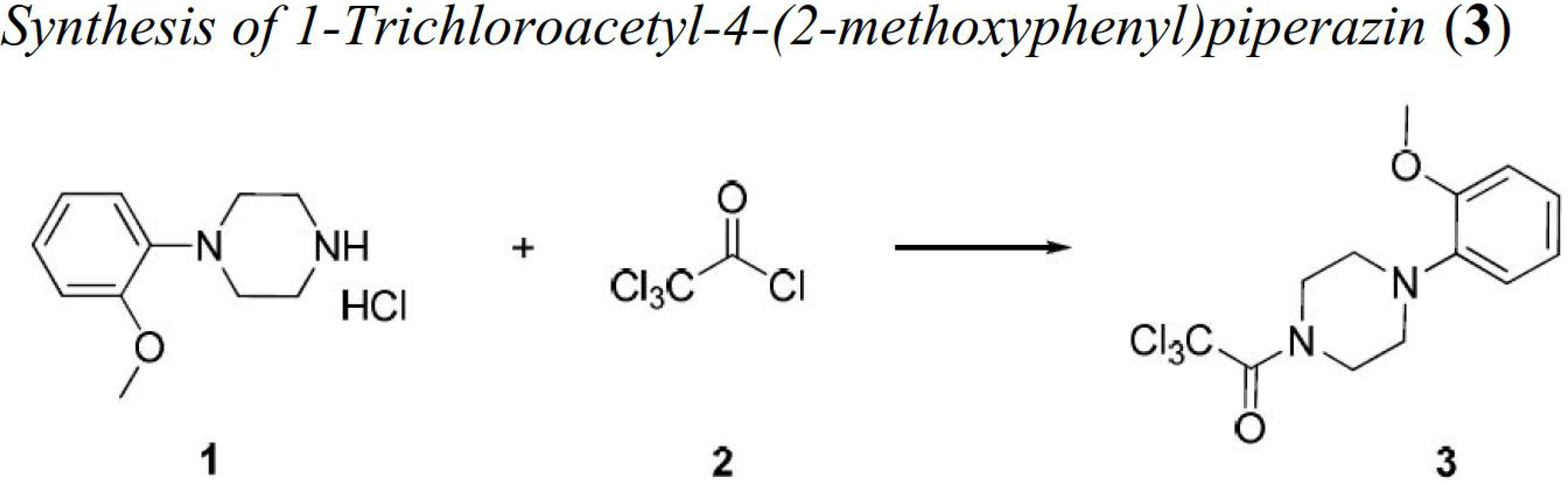

In a 2-L round-bottom flask equipped with thermometer, mechanical stirrer, dropping funnel and drying tube 1-(2-methoxyphenyl)piperazin hydrochloride (**1**) (91.00 g, 398 mmol) was suspended in dry dichloromethane (1000 mL). With well stirring and keeping the temperature below 5 °C trichloroacetyl chloride (**2**) (46.60 ml, 418 mmol, 1.05 equiv.) was added dropwise followed by the addition of N,N-diisopropylethylamine (146 ml, 855 mmol, 2.15 equiv.). The reaction mixture was let to warm up to room temperature and the stirring was continued for an additional hour, by that time the reaction was completed.

Water (ca. 800 ml) was added to the reaction mixture and after a few minutes the phases were separated. The aqueous part was extracted once with dichloromethane, the combined organic phases were washed with water and brine, dried over MgSO_4_ and evaporated. The crystalline residue was suspended in diisopropyl ether, filtered off and washed with the same solvent. The title product was dried in a vacuum desiccator over P_2_O_5_/KOH. Yield: 129.5 g of **3** (96%) as pale brown crystals.5 **LU-240 (3):** 1H NMR (200 MHz, *DMSO-d*6) δ ppm 6.86 – 7.02 (m, 11H), 3.70 – 4.10 (m, 7H); APCI MS *m/z* 337 [M + H]+; HPLC-MS >99.0% (AUC).

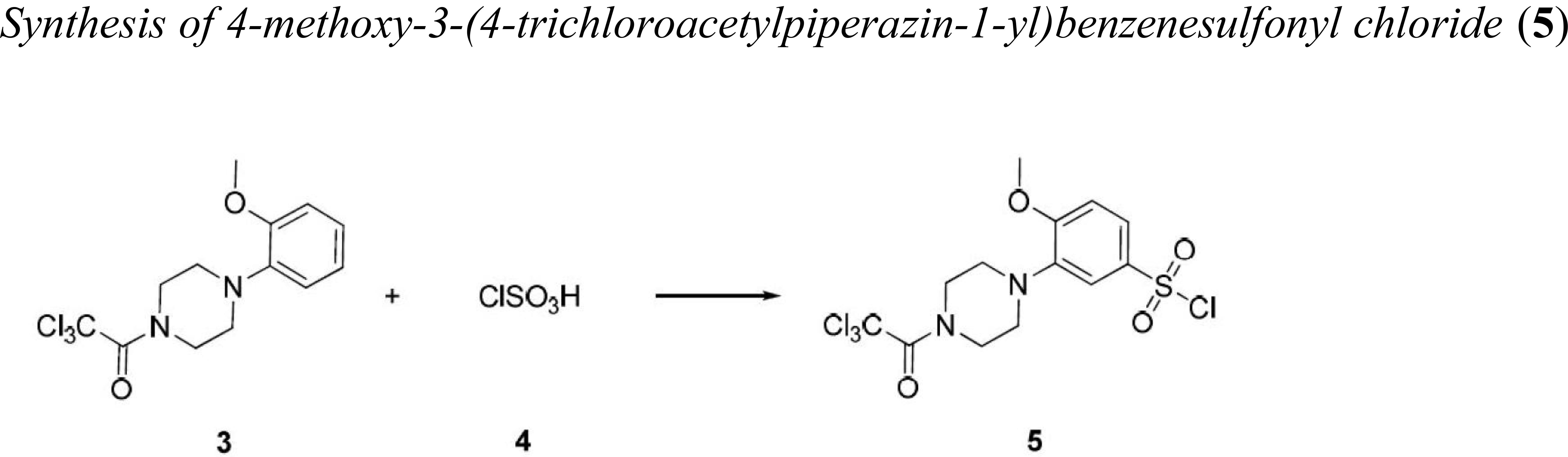

To a 2-L round-bottom flask equipped with thermometer, mechanical stirrer, dropping funnel and drying tube chlorosulfonic acid (**4**) (1.14 kg, 9.78 mol, 0.65 L) was charged. With well stirring and cooling to keep the temperature below 5 °C the solution of the piperazine derivative (**3**) (129.0 g; 384 mmol) in dry dichloromethane (500 ml) was added dropwise. The reaction mixture was let to warm up to room temperature and the stirring was continued for an additional hour.

To a five liter beaker, equipped with thermometer, mechanical stirrer and immersed into a saltice cooling mixture ca. 1 kilogram of ice was placed, and then the reaction mixture was slowly poured into it such a rate, that the temperature kept below 5 °C. The phases were separated and the aqueous part was extracted once with dichloromethane. The combined organic phase was washed with water and brine, dried over MgSO_4_ and evaporated to dryness. The crystalline residue was suspended in diisopropyl ether, filtered off and washed with the same solvent. Yield: 117.2 g of **5** (70%) as light brown crystals. **LU-238 (5):** ıH NMR (200 MHz, *DMSO-d*6) δ ppm 7.44 (s, 1H), 7.41 (d, *J* = 8.4 Hz, 1H), 7.03 (d, *J* = 8.4 Hz, 1H), 3.85 (s, 3H), 3.90 – 4.15 (m, 8H); APCI MS *m/z* 436 [M + H]+; HPLC-MS 95.0% (AUC).

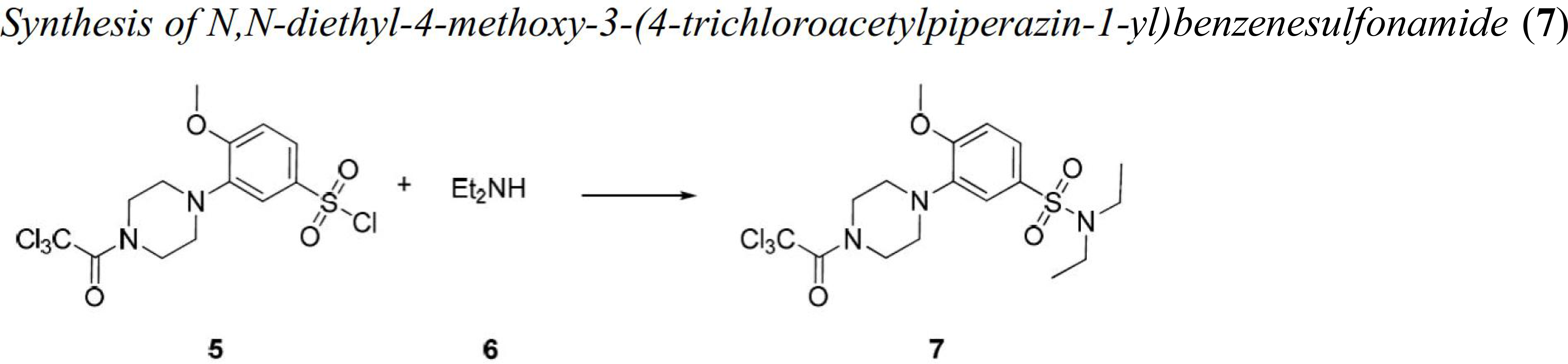

In a 1-L round-bottom flask equipped with thermometer, magnetic stirrer, dropping funnel and drying tube 21.30 ml of diethylamine (**6**) (15.10 g, 206 mmol, 3.00 equiv) was diluted with dry dichloromethane (300 mL). With well stirring and keeping the temperature below 5 °C the solution of sulfochloride derivative (**5**) (30.00 g; 69 mmol.) in dry dichloromethane (200 ml) was added dropwise. After the addition had taken place the reaction mixture was let to warm up to room temperature and the stirring was continued for additional 30 min, by that time the reaction has been completed.

The reaction mixture was diluted with water and the phases were separated. The organic phase was washed twice with water and once with brine, dried over MgSO_4_ and evaporated to dryness. The crystalline residue was suspended in diisopropyl ether, filtered off and washed with the same solvent. Yield: 31.30 g of 7 (96%) as light brown powder. **LU-247 (7):** ıH NMR (200 MHz, *DMSOd*6) δ ppm 7.44 (d, *J* = 8.4 Hz, 1H), 7.17 (s, 1H), 7.14 (d, *J* = 8.4 Hz, 1H), 3.70 – 4.10 (m, 11H), 3.11 (q, *J* = 7.2 Hz, 4H), 1.02 (t, *J* = 7.2 Hz, 6H); APCI MS *m/z* 472 [M + H]+; HPLC-MS 96.0% (AUC).

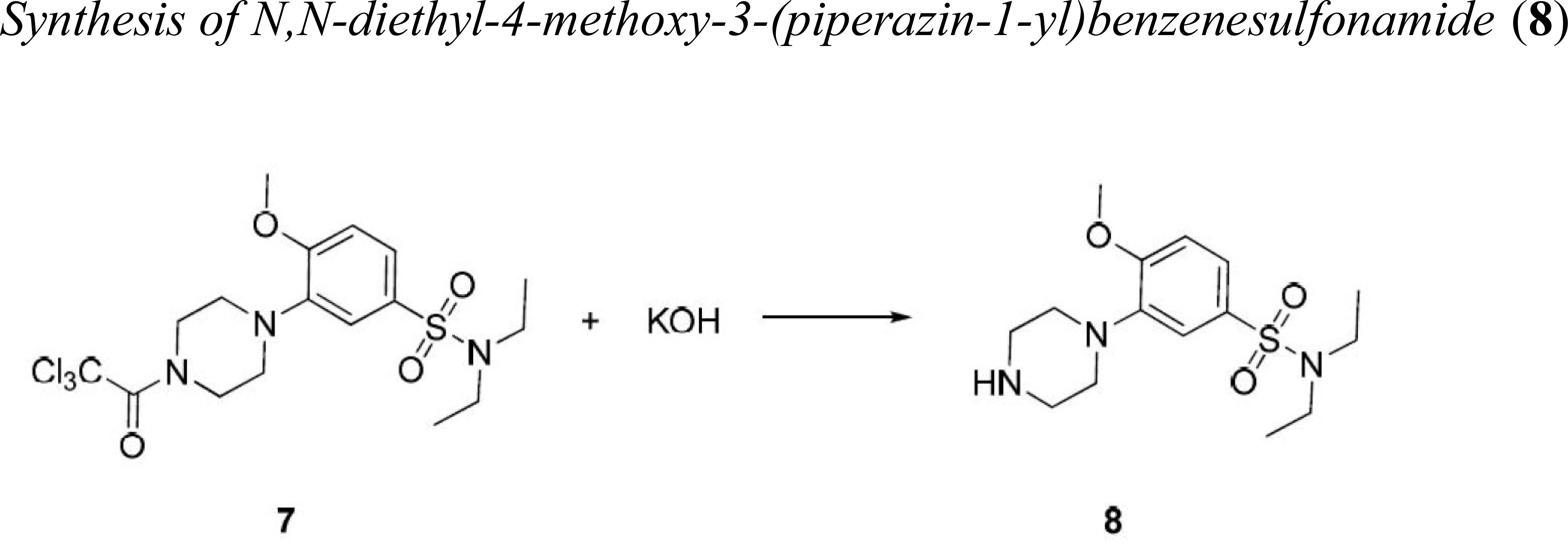

To a suspension of sulfonamide (**7**) (31.30 g, 66.2 mmol) in tetrahydrofuran (300 ml) was added the aqueous solution of potassium hydroxide (9.30 g, 166 mmol, 2.50 equiv., in 40 mL water) in one portion and the mixture was stirred at 40-50 °C for four hours. The same amount of potassium hydroxide (9.30 g, 166 mmol, 2.50 equiv) was dissolved in water (10 mL) and it was also added to the reaction mixture and the stirring was continued at 50 °C for another four hours then at room temperature overnight.

After completion of the reaction the solvent was evaporated, the aqueous residue was diluted with water and extracted with dichloromethane three times. The targeted piperazine derivative (**8**) was extracted from the organic phase three times with 3% HCl solution. The combined aqueous solution was cooled off, basified with 20% aqueous NaOH and extracted three times with dichloromethane. The combined organic solutions were washed with brine, dried over MgSO 4 and evaporated. The resulted yellow oil became crystalline on standing in refrigerator. It was suspended in a small amount of diisopropyl ether, filtered off and washed with the same solvent. Yield: 15.15 g of **8** (70%) as yellow crystalline. **LU-242 (8):** ıH NMR (200 MHz, *DMSO-d*6) δ ppm 7.34 – 7.48 (m, 1H), 7.05 – 7.20 (m, 2H), 3.86 (s, 3H), 2.90 – 3.20 (m, 12H), 2.61 (m, 1H), 1.02 (t, J = 7.2 Hz, 6H); APCI MS *m/z* 327 [M + H]+; HPLC-MS >99.0% (AUC).

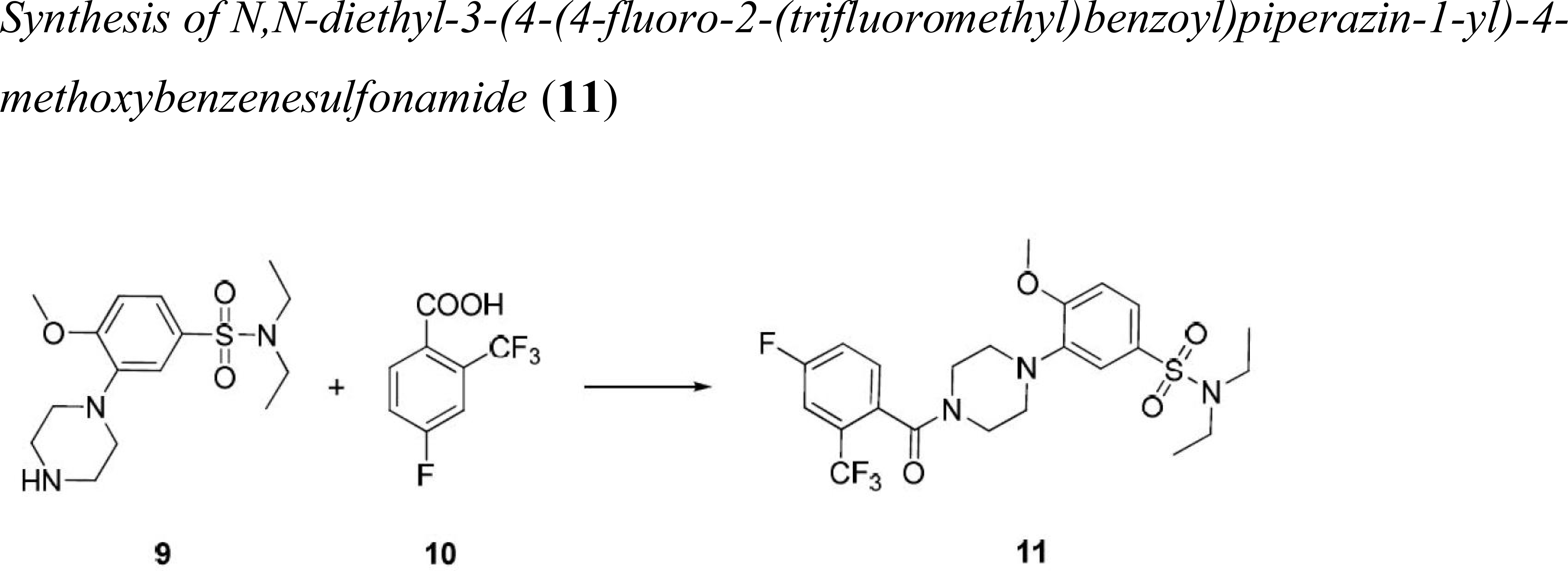

To a suspension of 4-fluoro-2-(trifluoromethyl)benzoic acid (**10**) (9.53 g, 46 mmol) in tetrahydrofuran (400 ml) was added 1,1’carbonyldiimidazole (8.17 g, 50 mmol, 1.10 equiv.) and the reaction mixture was stirred at room temperature for 2 hours. The solution of the piperazine derivative (**9**) (15.00 g, 46 mmol) in a minimal amount of tetrahydrofuran was added in one portion and the reaction mixture was stirred at 50 °C for 20 hours.

After the reaction had completed the solvent was evaporated. The residue was dissolved in dichloromethane and was washed with 3% aqueous HCl solution, water, saturated aqueous NaHCO_3_ solution and brine, dried over MgSO_4_, filtered and finally evaporated to dryness. This crude product was recrystallized from hot, 50% aqueous ethanol (500 ml) using norit. Yield: 12.91 g of **11** (54%) as white powder. **LU-246 (11):** 7.78 (d, *J* = 8.4 Hz, 1H), 7.61 – 7.69 (m, 2H), 7.08 – 7.18 (m, 2H), 3.87 (s, 3H), 3.70 – 3.82 (m, 4H), 2.75 – 3.35 (m, 8H), 1.01 (t, *J* = 7.2 Hz, 3H); APCI MS *m/z* 517 [M + H]+; HPLC-MS >99.0% (AUC).

Thin-layer chromatography (TLC) was performed using silica gel 60 F254 plates (Merck) and visualized by UV light (254 nm). Column chromatography was carried out on Biotage Horizon flash purification system using silica gel 20-40 unless otherwise specified. Proton nuclear magnetic resonance spectra were obtained on a Varian Unity 200 MHz instrument. For the calibration of spectra, solvent peak and tetramethylsilane signals were used. Spectra were performed at room temperature; the results are given in ppm (δ) with coupling constants and *J* values reported in hertz. The HPLC-MS analysis was performed on a Waters HPLC/MS (with 4-channel MUX interface) with a LiChroCART 30-4 Purospher STAR RP-18, endcapped, 3 m (Merck) column using a solvent gradient program.

##### BT13 NMR data

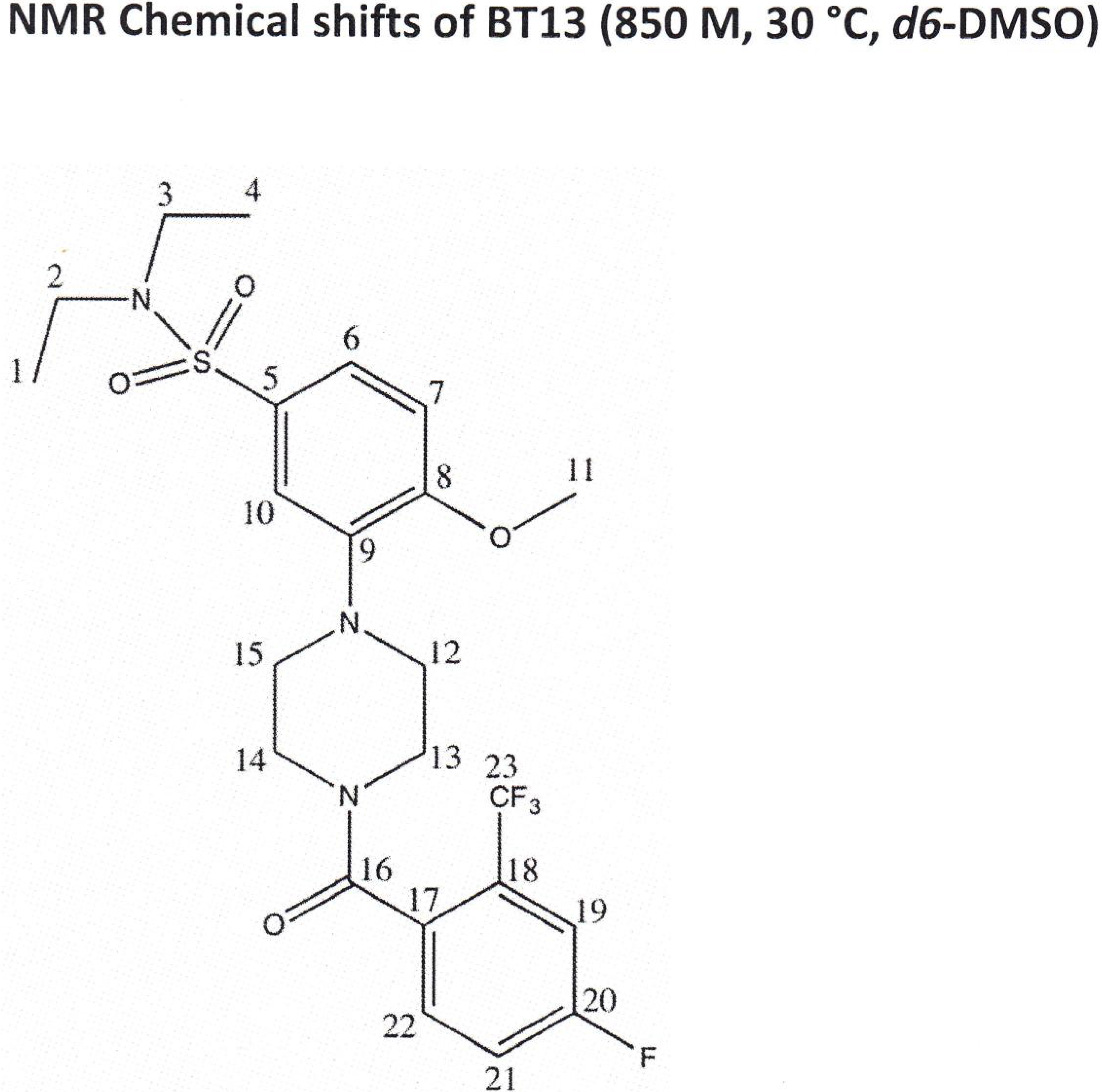

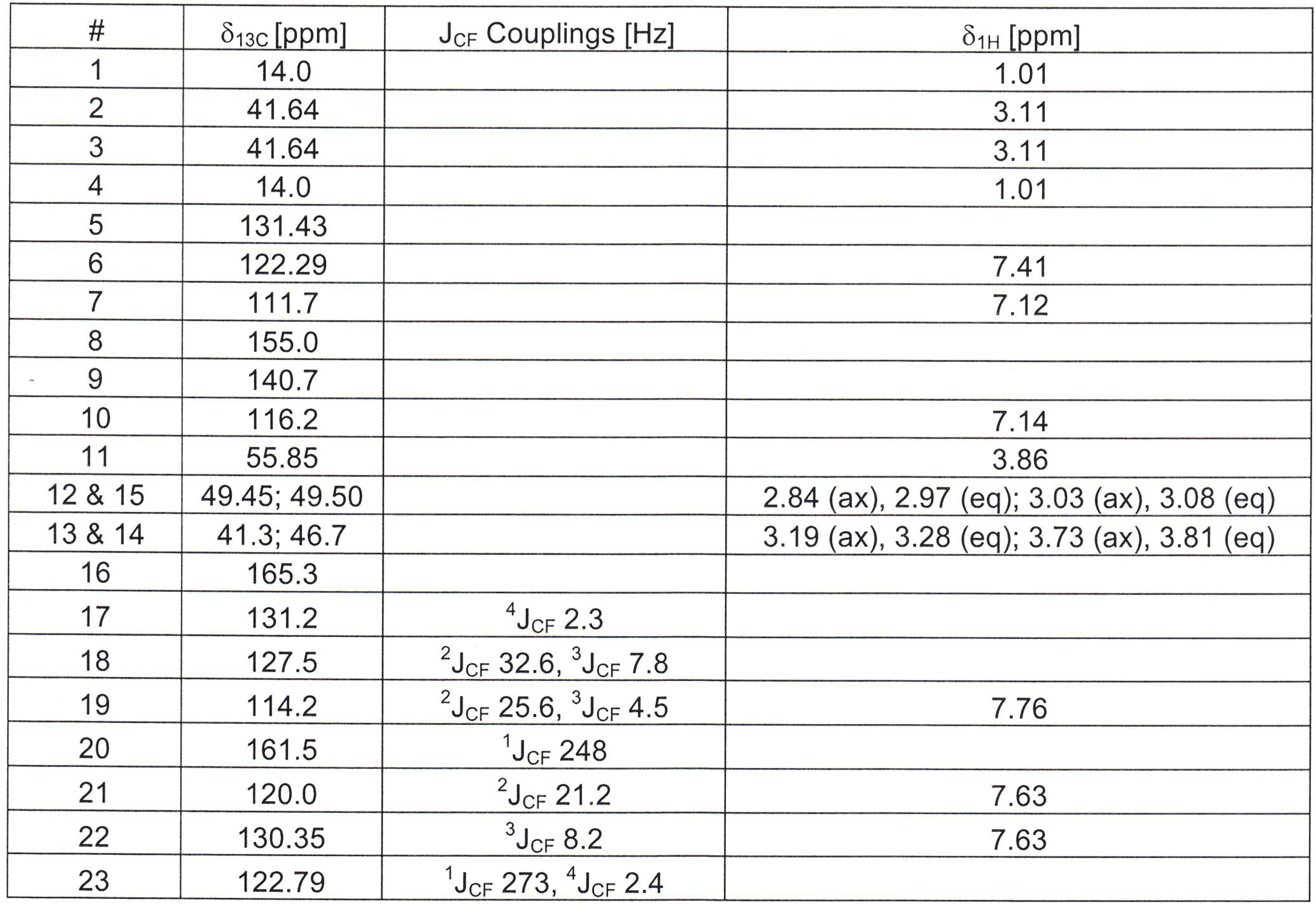

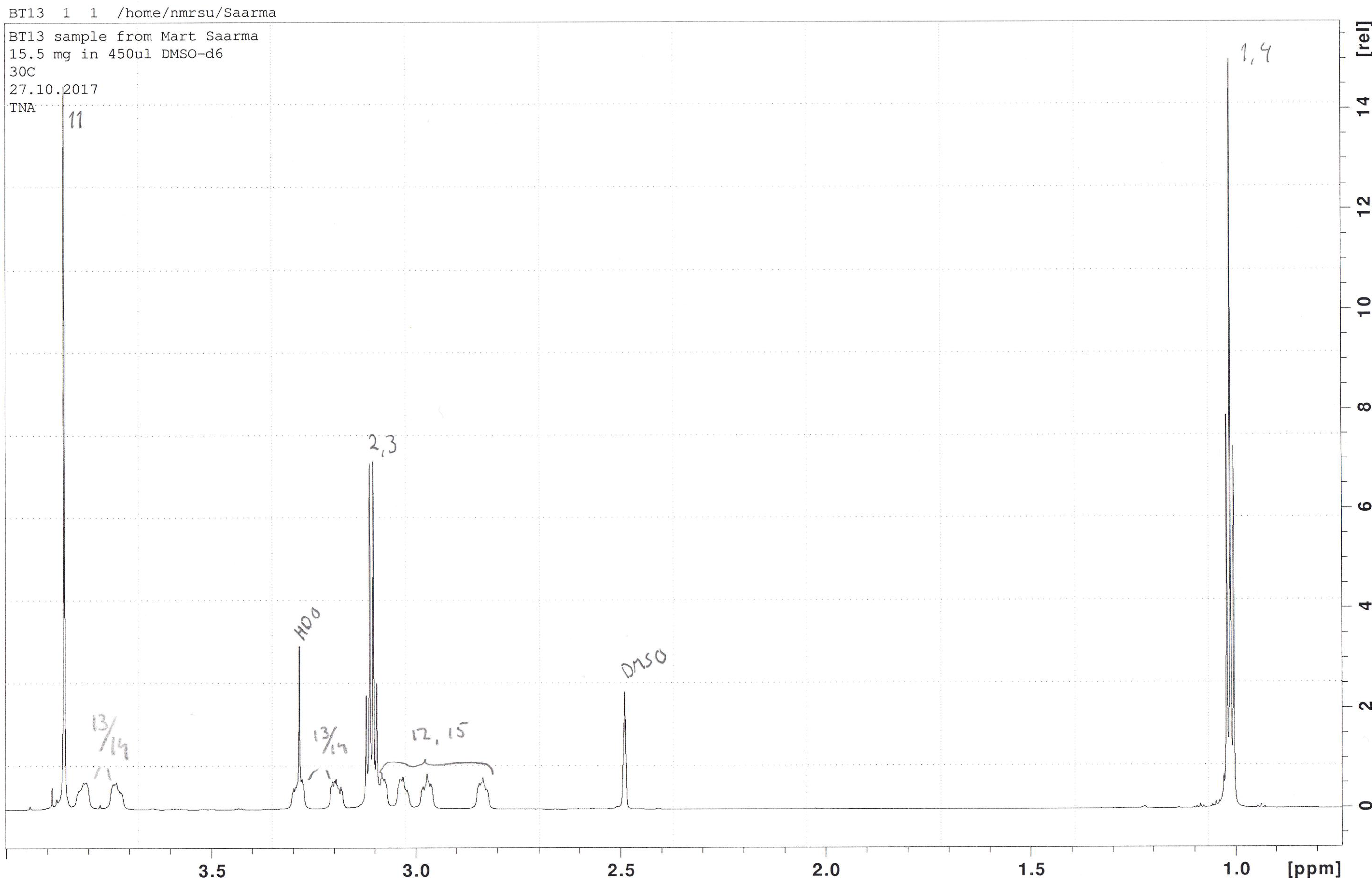

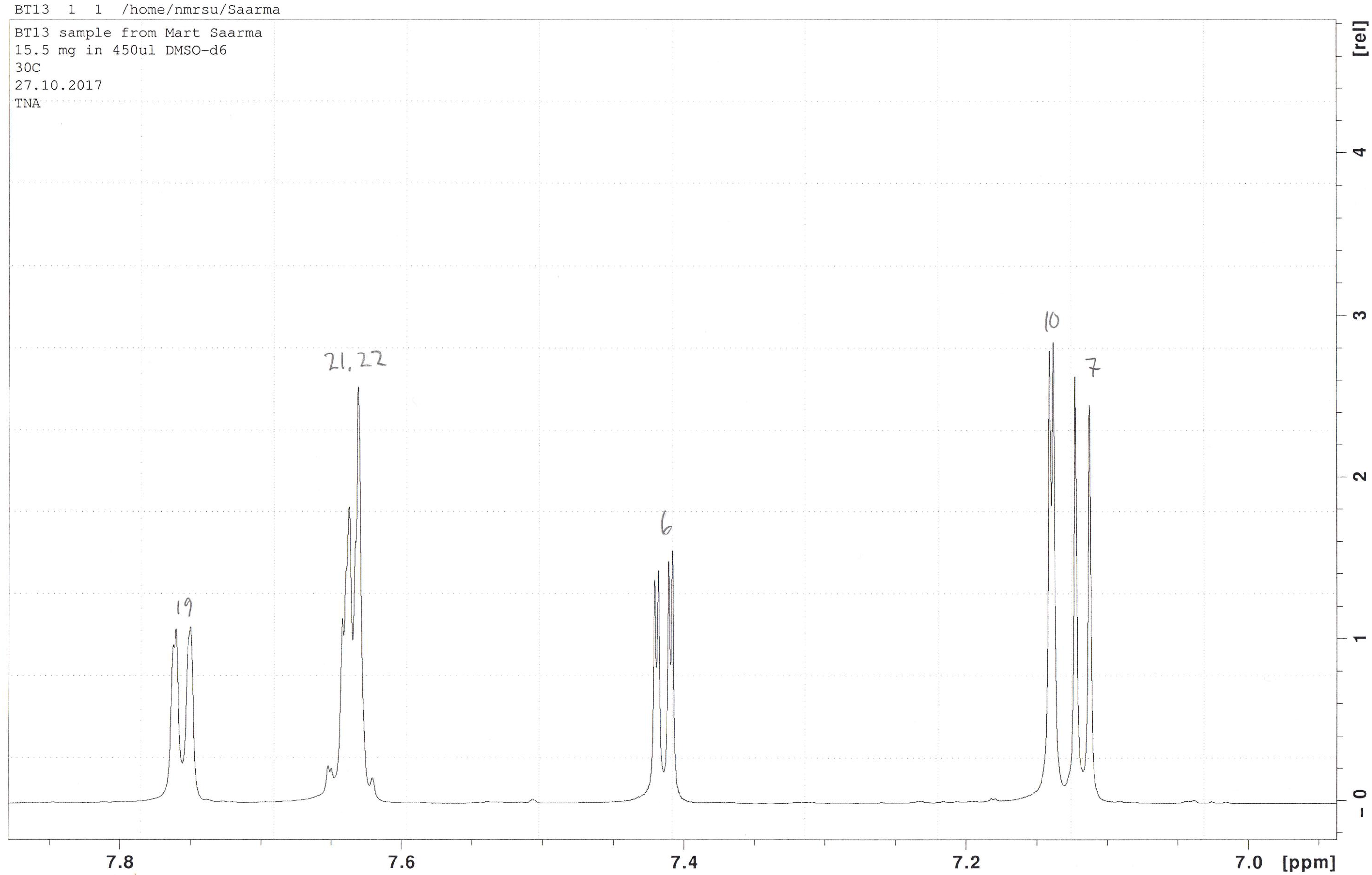

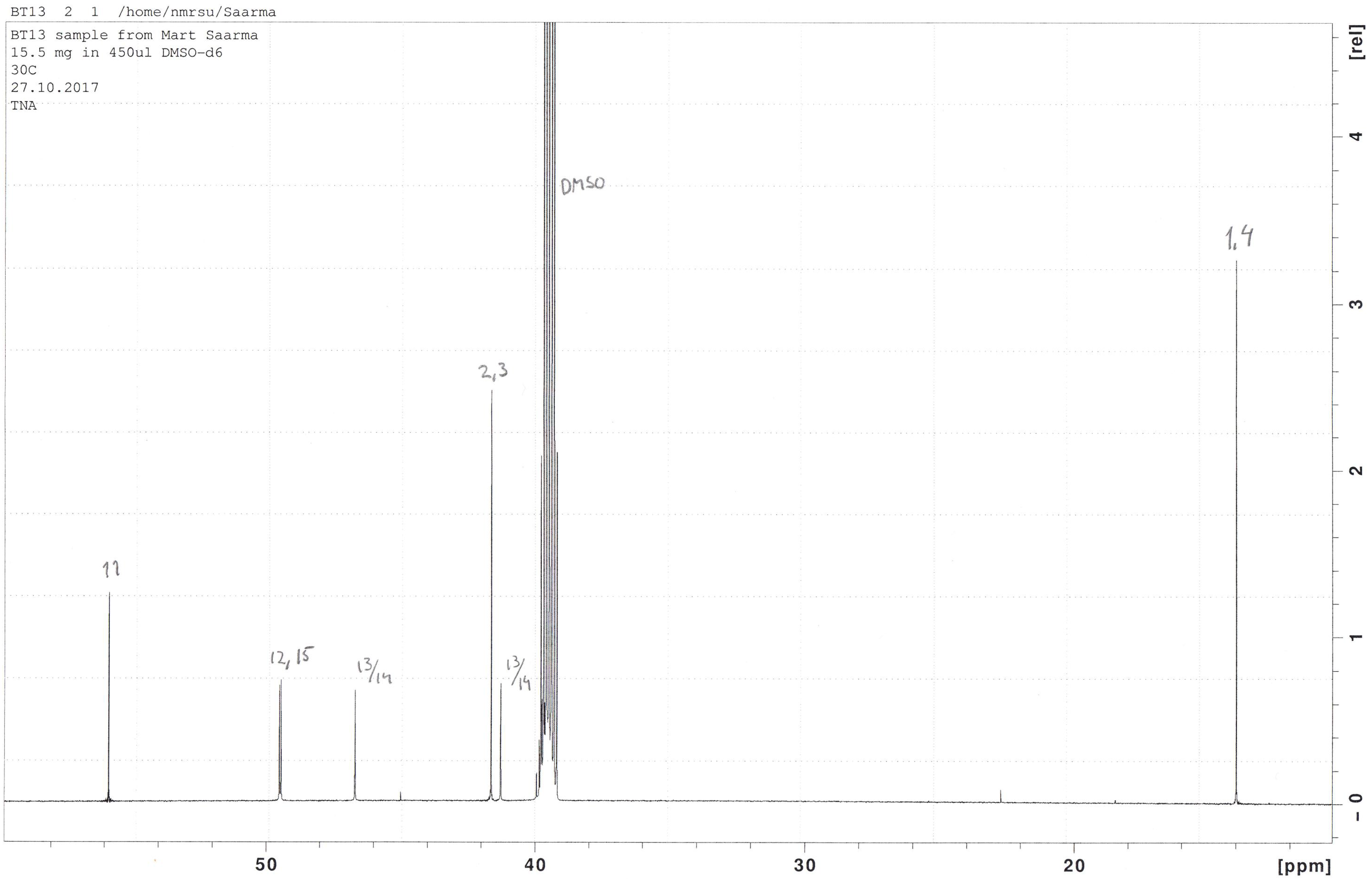

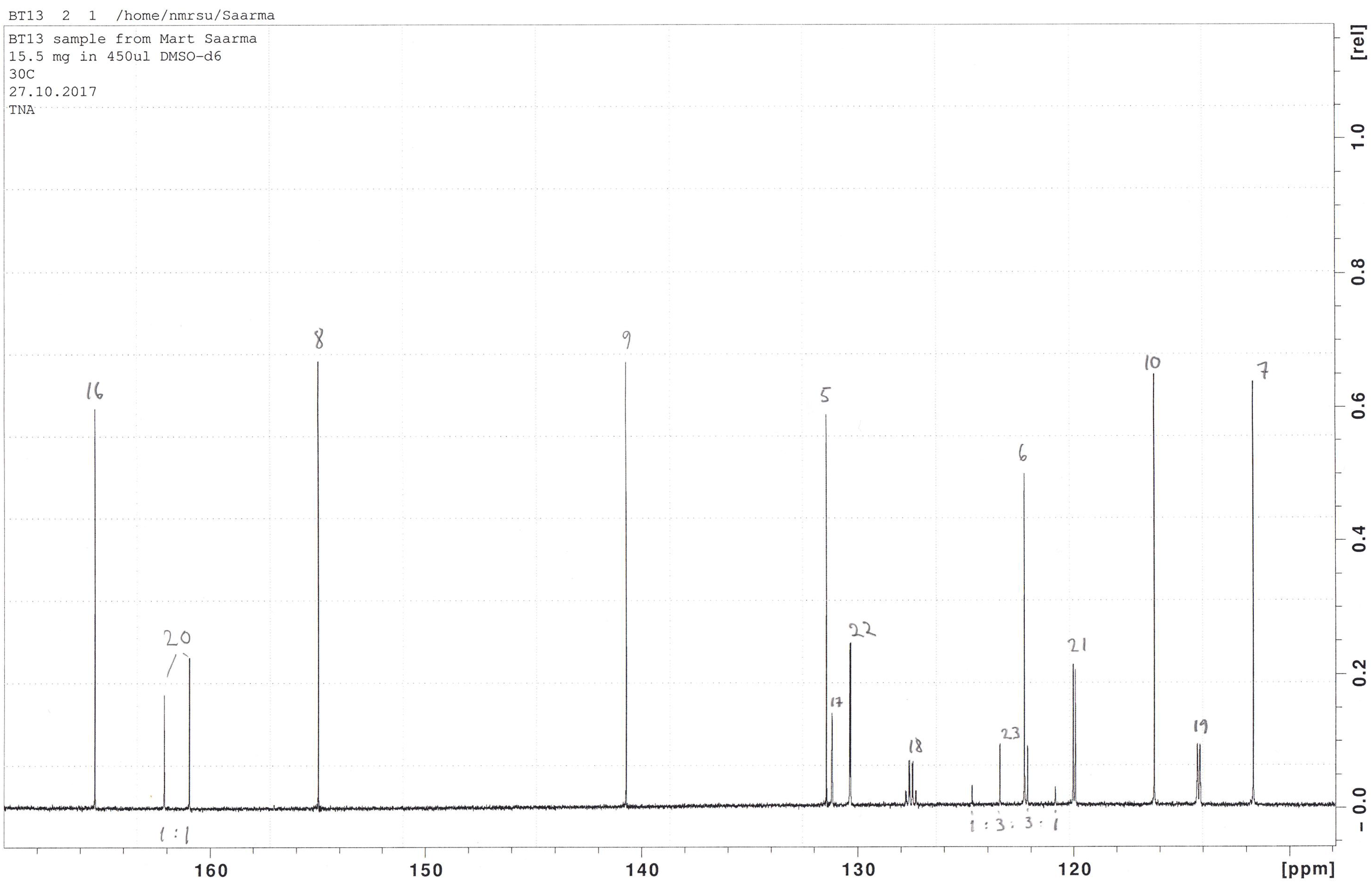

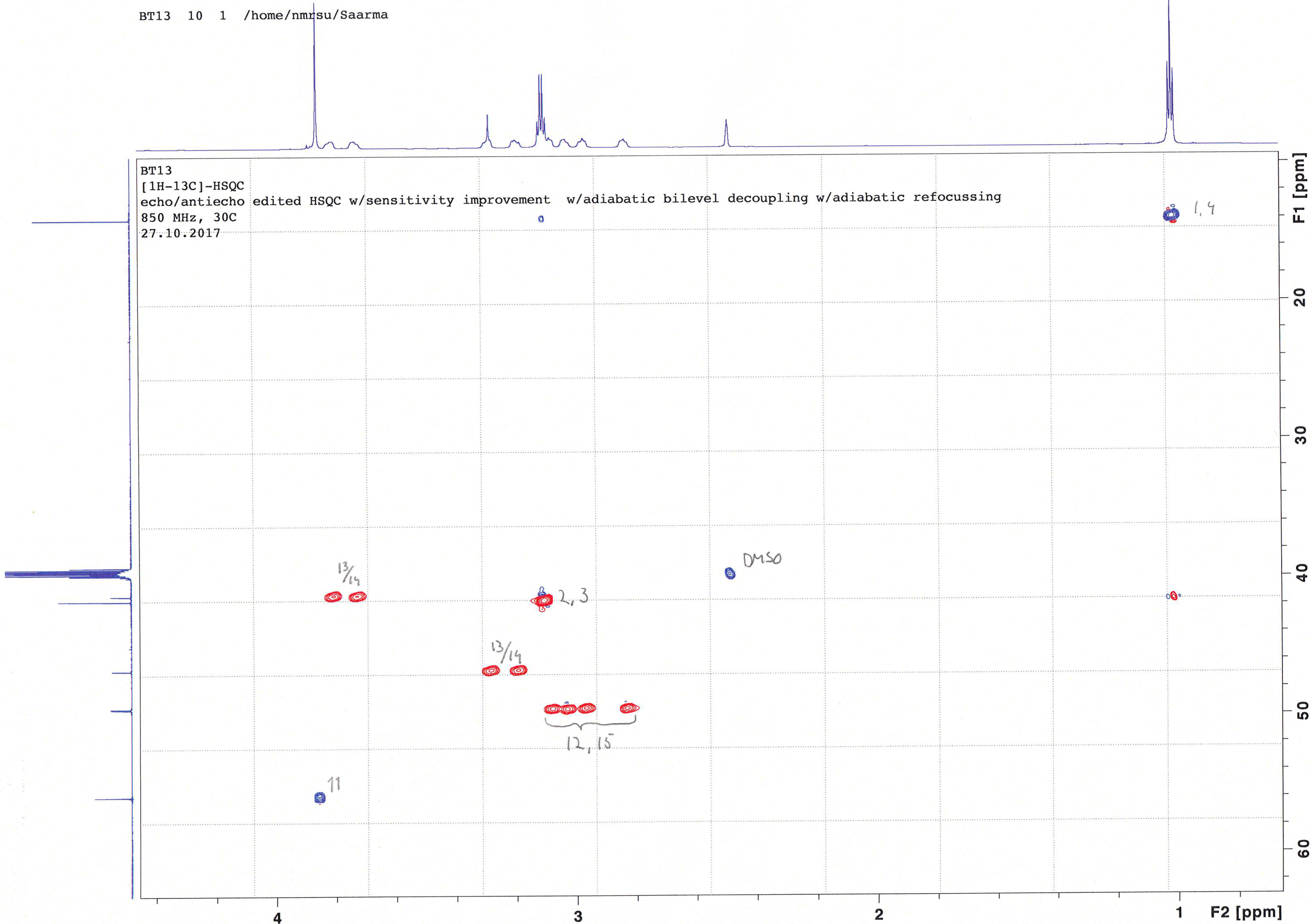

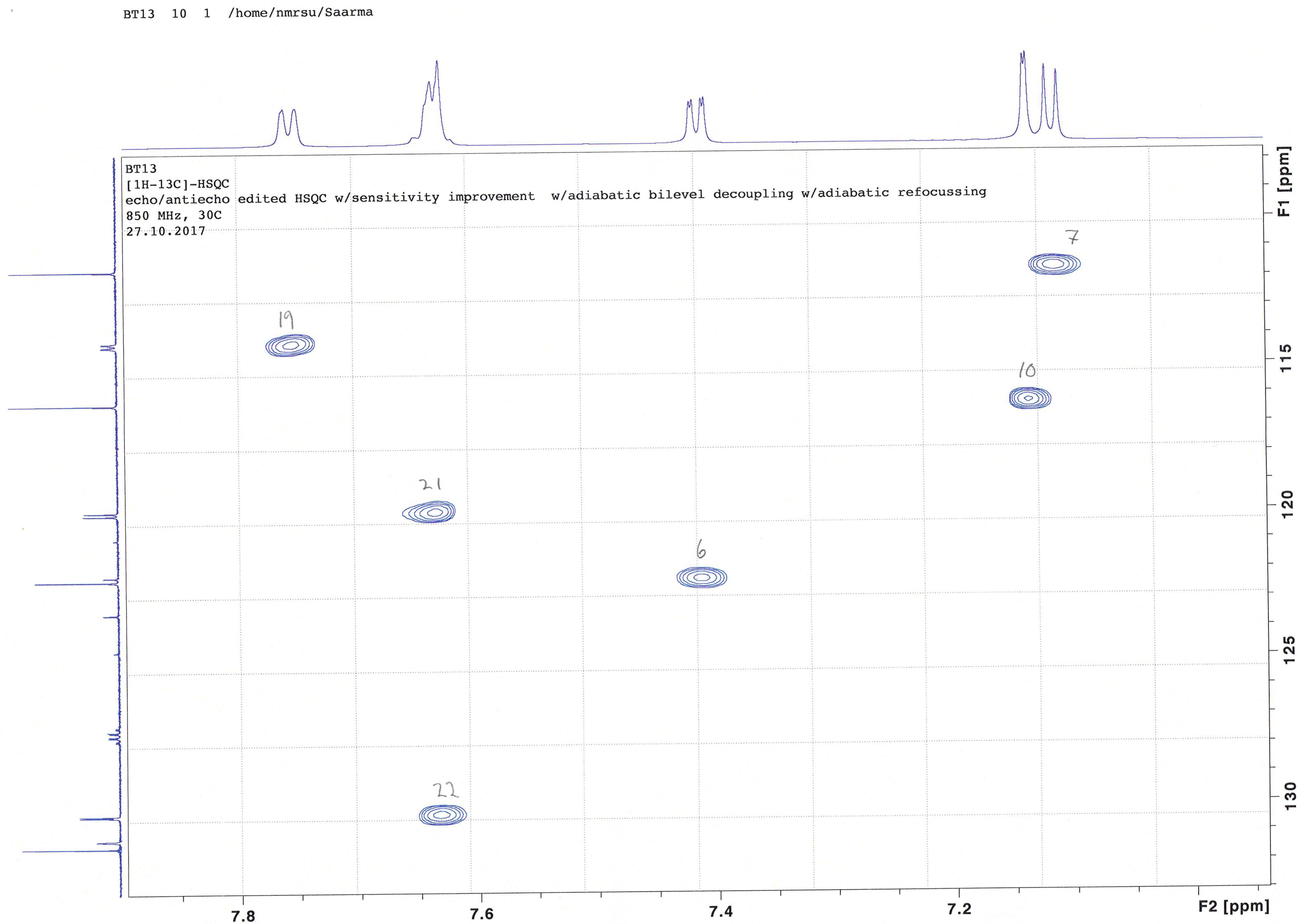

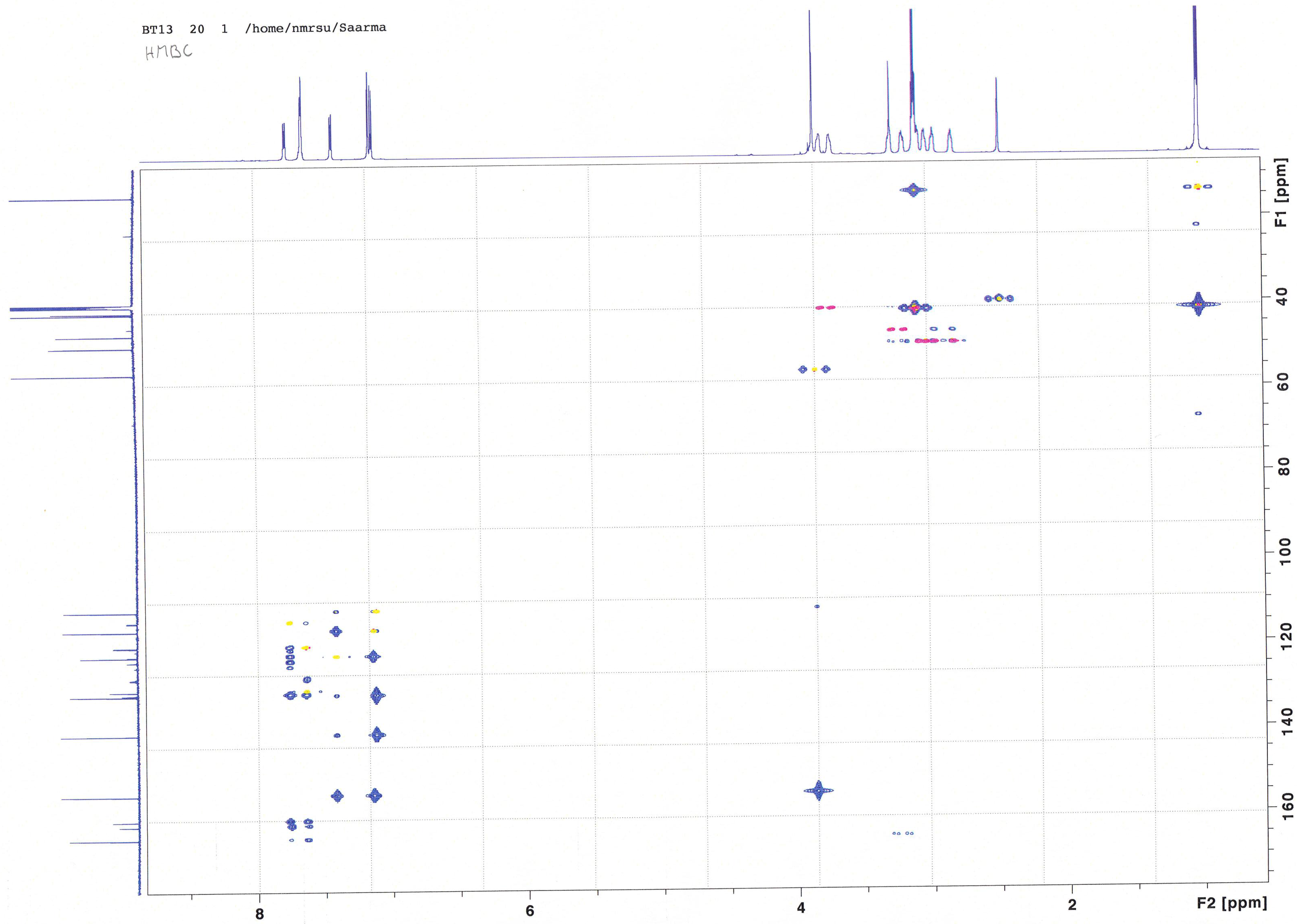

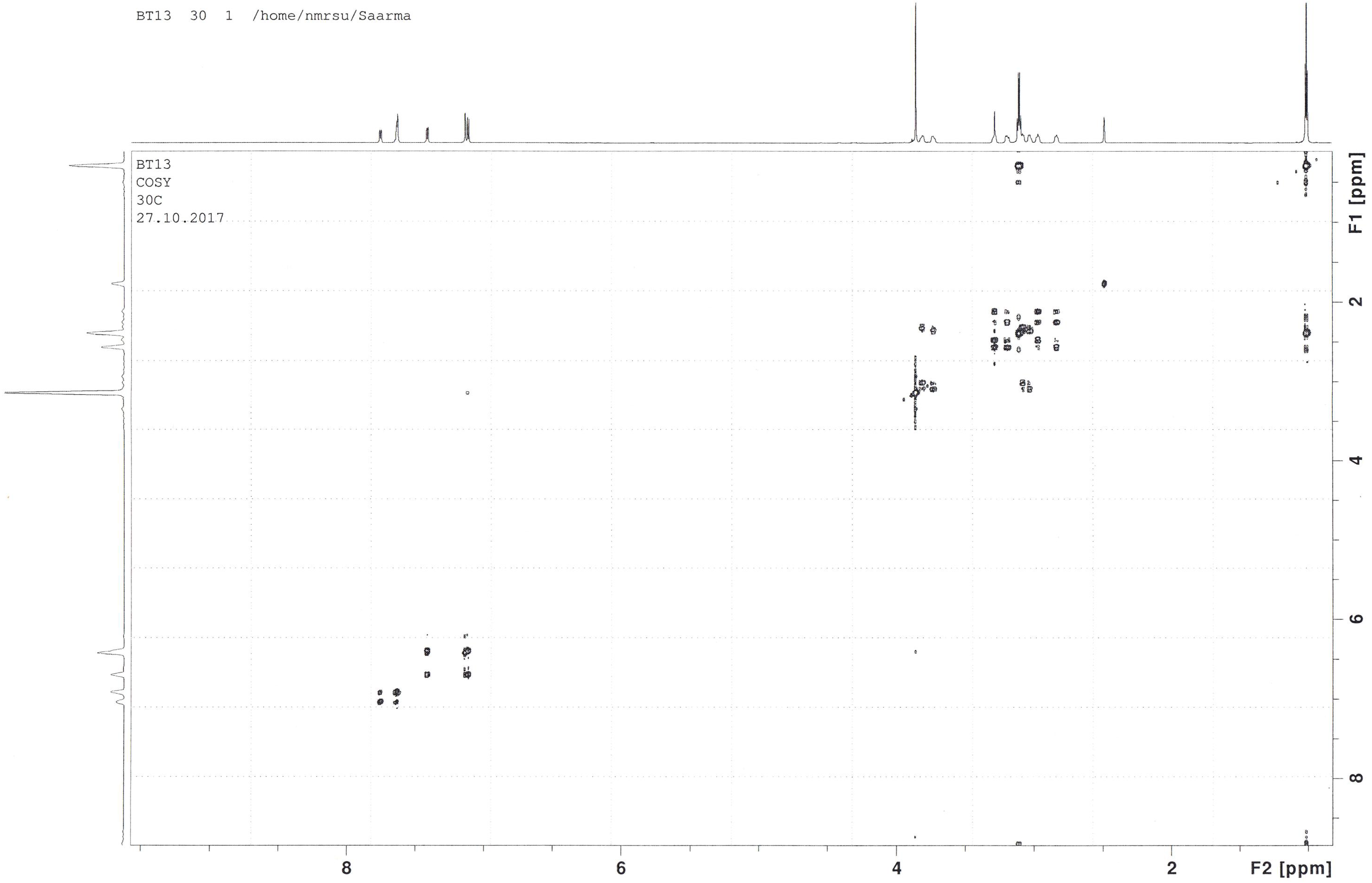

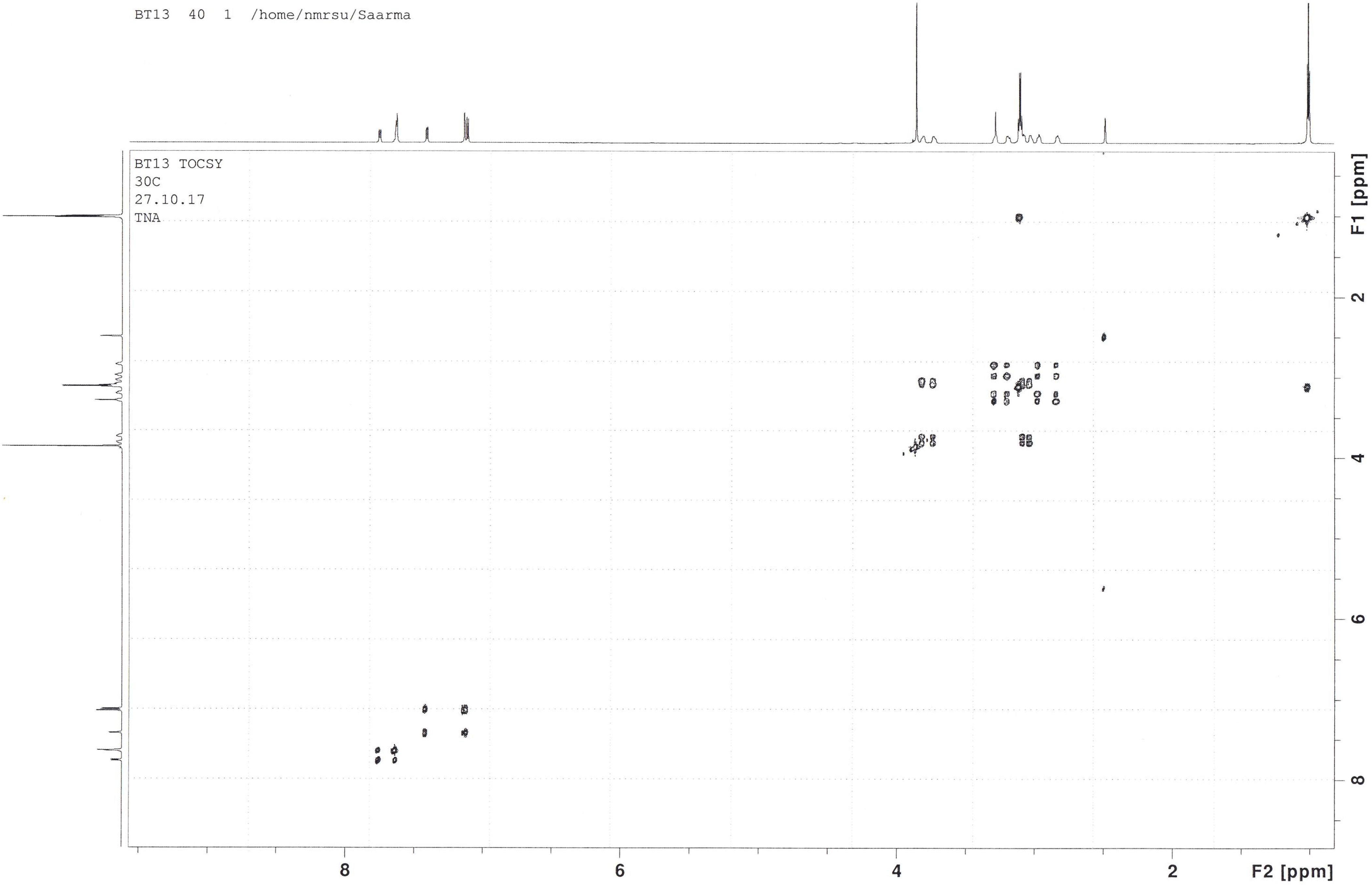

#### Experimental animals

Adult male mice C57Bl/6 (JRccHsd) (Harlan, the Netherlands) of 8–15 weeks of age weighing 19– 32 grams were used for brain microdialysis experiments and for immunohistochemistry of phosphorylated extracellular-signal-regulated kinases 1 and 2 (pERK) and phosphorylated ribosomal protein S6 (pS6) in striatum. E13.5 embryos of NMRI mice and RET knock-out mice (C57BL/6JOlaHsd) (Laboratory Animal Centre, University of Helsinki) were used for primary cultures of midbrain dopamine neurons. C57Bl/6 mice were housed individually or in groups of 2–10 animals per cage. Adult male Wistar rats (RccHan:WIST) (Harlan), weighing 240–435 grams at the start of the experiments were used for testing BT13 in the 6-OHDA model of Parkinson’s disease. Rats were housed in groups of 2–4 animals per cage, but after receiving intracerebral cannulas they were housed individually until removal of the cannulas. All animals were kept under 12:12 h light-dark cycle (lights on at 6 am) and ambient temperature in the animal facilities was 22 °C. Standard rodent chow (Harlan) and tap water were available *ad libitum*. Weight of the rats was monitored regularly, at least once per week, during the experiment. All experiments were carried out according to the European Community guidelines for the use of experimental animals and approved by the National Animal Experiment Board of Finland (license numbers: ESAVI/7551/04.10.07/2013, ESAVI/198/04.10.07/2014) for experiments with living animals and the Laboratory Animal Centre of the University of Helsinki (license number: KEK15-022) for collection of E13.5 embryos of NMRI mice.

### Genotyping

Mouse E13.5 embryos were genotyped using PCR for identification of embryos lacking RET expression as described previously (Schuchardt *et al.*, 1996). Tail snips from each embryo were collected in separate tubes and DNA was isolated using AccuStart II PCR Genotyping Kit (Quantabio, USA). Briefly, 100 l of DNA extraction reagent was added to each tube with a tail snip. The samples were boiled at 95°C for 30 minutes. After cooling to room temperature, equal volume of stabilization buffer was added to the tube. PCR reaction mixture was prepared by adding 1 l of DNA extract to mixture containing AccuStart II GEL Track PCR Supermix (2X), 400 nM primers P1-P4 (see below) and nuclease-free water. After the initial denaturation cycle (94°C, 2 minutes), DNA was amplified for 30 cycles in the following conditions: 94°C, 30 seconds; 60°C, 30 seconds; 72°C, 30 seconds. The sequences of the primers used in the reactions were: P1, 5′-TGGGAGAAGGCGAGTTTGGAAA-3′; P2, 5′-TTCAGGAACACTGGCTACCATG-3′ for wild allele and P3, 5′-AGAGGCTATTCGGCTATGACTG-3′; P4, 5′-CCTGATCGACAAGACCGGCTTC-3′ for mutant allele.

### Luciferase assay

Luciferase assay was performed as described earlier (Sidorova *et al.*, 2010, 2017). Reporter cells (200 000 cells/ml) were plated to 96-well tissue culture plates (100 μl/well) a day before stimulation with BT13 or GDNF in DMEM, 10% fetal bovine serum (FBS), 15 mM HEPES pH 7.2, 1% DMSO, 100 g/ml normocin. The next day 100 μl of 2xGDNF or 2xBT13 solutions in DMEM, 15 mM HEPES, pH 7.2 and 1% DMSO were added to the reporter cells. The cells were cultured in the presence of the tested substances for 24 hours to produce luciferase. Luciferase activity was measured using neolite reagent (PerkinElmer, USA) according to the manufacturer’s instruction. The luminescence was detected using FLUOstar OPTIMA microplate reader (BMG LABTECH). Compounds were tested in quadruplicates in 3 independent experiments.

### Cell culture, cell transfection, treatments and sample preparation for phosphorylation assays

MG87RET cells were cultured overnight in 6-well plates and transfected with hGFRα1, hGFRα2 and GFP-expressing plasmid using Lipofectamine 2000 (Invitrogen), as described by the manufacturer. The next day, cells were starved for 4 hours using starvation medium (DMEM, 15 mM HEPES pH 7.2, 1% DMSO). The cells were stimulated by different concentrations of BT13 (10, 25, 50, 100 μM) and GDNF (250 ng/ml) or NRTN (250 ng/ml) dissolved in starvation medium for 15 minutes. Then the cells were washed with ice-cold PBS (phosphate buffered saline, pH 7.4; 137 mM NaCl, 2.7 mM KCl, 10 mM PO_4_^3-^) containing 1 mM Na_3_VO_4_ and lysed with 500μl RIPA-modified buffer (50 mM Tris-HCl, pH 7.4, 150 mM NaCl, 1 mM EDTA, 1% NP-40, 1% TX-100, 10% glycerol, EDTA-free protease inhibitor cocktail (Roche, Switzerland), 1 mM Na_3_VO_4_, 6 mM sodium deoxycholate, 1 mM PMSF) per well.

### Western blotting-based ERK and AKT phosphorylation assay

Levels of pERK and pAKT in immortalized cells were analyzed as described previously (Sidorova *et al.*, 2017). Cell lysate (100 μl) (see above) was collected and mixed with the same amount of 2xLaemmli buffer. Samples were boiled for 10 minutes and proteins were resolved on a 12% sodium dodecyl sulfate–polyacrylamide gel SDS-PAAG. After electrophoresis, proteins were electro-transferred onto nitrocellulose membranes. Lower part of the membranes containing ERK among others proteins was blocked with 10% skimmed milk in TBS-T (TBS (50 mM Tris-HCl, pH 7.4, 150 mM NaCl) with 0.15% of Tween 20) whereas the upper part containing AKT among other proteins was blocked with 10% BSA in TBS-T for 10 minutes and probed with E4 pERK (1:1000 in 3% skimmed milk in TBS-T; Santa Cruz Biotechnology Cat# sc-7383, RRID:AB_627545) and pAKT (1:500 in 3% BSA in TBS-T; Cell Signaling Technology Cat# 9271, RRID:AB_329825) antibodies, respectively, overnight at 4°C. The next day, both membranes were washed 3 times for 10 minutes with TBS-T. ERK-containing membrane was incubated with horseradish peroxidase (HRP)-conjugated anti-mouse secondary antibody (DAKO, Cat# P0447), diluted 1:3000 in 3% skimmed milk in TBS-T, and AKT-containing membrane was incubated with HRP-conjugated anti-rabbit antibody (GE Healthcare Cat# NA934, RRID:AB_772206), diluted 1:3000 in 3% BSA in TBS-T for 1 hour at room temperature. The membranes were washed again 4 times for 15 minutes with TBS-T. Bands were visualized with ECL Plus Western Blotting Substrate (Pierce, USA) or SuperSignal West Femto Maximum Sensitivity Substrate (Pierce) using Luminescent Image Analyzer LAS3000 (Fuji-Film, Japan). Membranes were stripped and re-probed with GAPDH antibody (1:4000, Millipore Cat# MAB374, RRID:AB_2107445) for 1 hour at room temperature to confirm equal loading of the proteins to the gels. Membranes were washed and incubated with HRP-conjugated anti-mouse secondary antibody for 1 hour and visualized according to the above-mentioned procedure.

### Analysis of RET phosphorylation by Western blotting

The level of RET phosphorylation in the cells was analyzed by Western blotting as described previously (Leppanen *et al.*, 2004). Cell lysates prepared as described above were centrifuged (5000 rpm, 5 min, 4ºC) to precipitate cell debris. RET was immunoprecipitated from the supernatant with 2 μg/ml anti-RET C-20 antibody (Santa Cruz Biotechnology Cat# sc-1290, RRID:AB_631316) and Dynabeads Protein G (Thermo Fisher Scientific) overnight at 4ºC. The next day, the beads were washed 3 times with TBS with 1% Triton X-100. Immunoprecipitated proteins were eluted by adding 50 μl of 2xLaemmli buffer, resolved on 7.5% SDS-PAAG and transferred to nitrocellulose membrane. The membrane was blocked with 10% skimmed milk in TBS-T for 10 minutes and probed overnight with phospho-tyrosine-specific antibody (1:1500 in 3% skimmed milk in TBS-T; Merck Millipore Cat# 05-321, RRID:AB_309678). The membrane was washed 3 times for 10 minutes with TBS-T and incubated with HRP-conjugated anti-mouse secondary antibody (1:3000 in 3% skimmed milk in TBS-T) for 1 hour at room temperature. The membrane was washed again with TBS-T 4 times for 15 minutes. Bands were visualized with with ECL Plus Western Blotting Substrate or SuperSignal West Femto Maximum Sensitivity Substrate using Luminescent Image Analyzer LAS3000. Loading control was confirmed by re-probing the membrane with anti-RET C-20 antibody (1:500) for 1 hour at room temperature after stripping. The membrane was washed and incubated with HRP-conjugated anti-goat secondary antibody (1:1500, DAKO, Cat# P0449) for 1 hour and visualized according to the above-mentioned procedure.

### RET-ELISA assay

To quantify the level of phosphorylated RET in immortalized cells we performed phospho-RET-ELISA assays (Parkash *et al.*, 2008). Assay plates (OptiPlate 96 F HB, Wallac) were coated with anti-RET C-20 antibody (1 μg/ml in TBS overnight), washed 4 times with PBS, blocked with blocking solution (5% BSA in TBS), 3 hours at room temperature and then washed with RIPA modified buffer. Cell lysates prepared as described above were added to the plates (70 μl/well) and incubated overnight at 4°C on a shaker. The plates were washed 3 times with washing buffer (TBS, 1% Triton X-100, 2% glycerol) and probed with phospho-tyrosine-specific antibody (1:1000 in binding buffer (TBS, 1% Triton X-100, 2% glycerol, 2%BSA)) for 1 hour at room temperature. The plates were washed 3 times with washing buffer, incubated with HRP-conjugated anti-mouse secondary antibody (1:3000 in binding buffer) for 1 hour at room temperature and washed once more 3 times with washing buffer. To detect the signal, pre-warmed SuperSignal West Femto Maximum Sensitivity Substrate (100 μl/well) was applied to the plates and luminescence was counted using FLUOstar OPTIMA microplate reader (BMG LABTECH).

### Survival assay for naïve dopamine neurons

#### Plate preparation

Plates were prepared as described previously (Planken *et al.*, 2010) with small modifications. A 96-well plate was pre-coated with poly-DL-ornithine, (0.5 mg/ml in 0.15 M borate buffer, pH 8.7, Sigma-Aldrich, Cat# P8638), overnight at 4ºC. The plate was washed 3 times with PBS and a micro-island was created in each well by circling at the edge of the well with the tip of a suction pump. The plate was dried for 1 hour in laminar flow hood and stored at 4°C before use.

#### Dissection and plating of dopamine neurons

Dopamine neurons were isolated from E13.5 embryos of NMRI mice. The midbrain floors were dissected in Dulbecco’s medium (0.1 g/l MgCl_2_•6H_2_O, 0.1 g/l CaCl_2_, 8 g/l NaCl, 0.2 g/l KCl, 1.4 g/l Na_2_HPO_4_•2H_2_O, 0.2 g/l KH_2_PO_4_) containing 2% BSA under a dissection microscope (Olympus SZX10 Stereo Microscope), washed 3 times with calcium and magnesium-free Hank’s Balanced Salt Solution (HBSS: Gibco, Life Technologies) and incubated in 5 mg/ml trypsin solution in HBSS for 20 minutes at 37°C. Enzymatic activity of trypsin was blocked by adding equal volume of FBS, containing 0.1 mg/ml of DNase I (Roche, Cat# 11284932001). Cells were triturated with a siliconized glass Pasteur pipette to get single cell suspensions and centrifuged at 1000 rpm for 5 minutes. The pellets were washed 3 times with primary neuron culture medium [(Dulbecco’s MEM/Nut mix F12 (Invitrogen/Gibco, Cat# 21331-020), 1×N_2_ serum supplement (Invitrogen/Gibco, Cat# 17502-048), 33 mM D-glucose (Cat # G-8769, Sigma-Aldrich), 0.5 mM L-glutamine (Invitrogen/Gibco, Cat# 25030-032), and 100 μg/ml Primocin (InvivoGen, Cat# ant-pm-2)] to remove the traces of serum. The washed pellets were resuspended in 150–200 μl of the primary neuron culture medium. The cells were counted using a TC20 automated cell counter (BIO-RAD). About 30000 cells were plated per well of previously prepared plates and cultured at 37°C.

Different concentrations of BT13 (0.01, 0.1, 1, 10 μM) and GDNF (Icosagen, 10 ng/ml) were dissolved in primary neuron culture medium containing 1% DMSO and applied to the wells within 1 hour post plating. The cells were incubated for 5 days and half of the culture media were replaced with fresh portions 2.5 days post plating.

#### Tyrosine hydroxylase immunocytochemistry

Dopamine neurons were visualized by immunocytochemical staining with an antibody against tyrosine hydroxylase (TH), the key enzyme of dopamine synthesis. After 5 days of culturing, cells were fixed with 4% paraformaldehyde (PFA) for 20 minutes and washed 3 times with PBS followed by permeabilization with 0.2% Triton X-100 in PBS for 15 minutes. Unspecific binding was blocked by incubating cells with a blocking solution (5% horse serum in 0.2% Triton X-100 in PBS) for 1 hour. Anti-TH antibody (1:500 in blocking solution, Millipore Cat# MAB318, RRID:AB_2201528) was applied to the cells and incubated overnight at 4°C. Cells were washed 3 times with PBS and incubated with Alexa Fluor 647 conjugated anti-mouse secondary antibody (1:500 in blocking solution, Thermo Fisher Scientific Cat# A-31571, RRID:AB_162542) for 1 hour at room temperature. Cells were washed with PBS to remove unbound antibody and nuclei were stained with 0.2 μg/ml DAPI (4’, 6-diamidino-2-phenylindole) in PBS for 10 minutes at room temperature. Finally, cells were washed 3 times with PBS and kept in PBS until imaging.

#### Image acquisition and analysis

Cells were imaged by CellInsight (CX51110, ThermoFisher Scientific) CX5 High Content Screening (HCS) with 20×magnification. Images were analyzed using CellProfiler image analysis software (Carpenter *et al.*, 2006).

### Survival of MPP^+^ challenged dopamine neurons

The survival analysis of MPP^+^ challenged wild-type dopamine neurons was performed by Neuron Experts company (http://www.neuronexperts.com/) using previously described method (Schinelli *et al.*, 1988). Ventral portion of the mesencephalic flexure was dissected from the brains of rat E15 embryos and trypsinized for 20 minutes at 37°C in 1% trypsin-EDTA (Invitrogen) diluted in Ca^2+^ and Mg^2+^-free PBS. The reaction was stopped by addition of DMEM containing 0.1 mg/ml DNAase I (Roche) and 10% FBS. Cells were dissociated by trituration, precipitated by centrifugation, plated on poly-L-lysine precoated 96-well plates (69 000 cells/well) and cultured in Neurobasal medium (Invitrogen) supplemented with 2% B27 (Invitrogen), 0.2 mM L-glutamine and 1% Penicillin-Streptomycin. Half of the culture medium was changed every two days with fresh medium. On 6^th^ day in vitro (6^th^ DIV) the culture medium was replaced by the fresh media containing 16 μM MPP^+^ and tested substances or DMSO as a negative control. Brain-derived neurotrophic factor (BDNF) (10 ng/ml) that is known to promote the survival of cultured dopamine neurons (Hyman *et al.*, 1991) was used as a positive control. After 48 hours (8^th^ DIV) cells were fixed with 4% PFA. Dopamine neurons were labelled using antibody against TH (Sigma-Aldrich Cat# T1299, RRID:AB_477560) and Alexa Fluor 488 conjugated goat anti-mouse antibody (Molecular Probes Cat# A-11017, RRID:AB_143160). Nuclei were labelled with Hoechst dye. For each condition, 2×10 pictures per well were taken using InCell AnalyzerTM 1000 (Amersham Biosciences, UK) with 10x magnification. All images were taken using the same conditions. The number of TH-positive neurons was analysed using InCell AnalyzerTM 1000 3.2. Workstation software.

### *In vivo* microdialysis

For *in vivo* microdialysis 2 months old C57BL/6J male mice were used. Each animal underwent 2 dialyses with 2 days washout period between them. A microdialysis guide cannula (MAB 4.1, AgnTho’s, AB, Sweden) was inserted into the dorsal striatum (AP = +0.6; ML = +1.8; DV = −2.2 relative to the bregma, according to the mouse brain atlas (Paxinos and Franklin, 2001) under isoflurane (Vetflurane 1000 mg/g, Virbac, France) anaesthesia and attached to the skull by two stainless steel screws and dental cement (Aqualox; Voco, Germany). Lidocaine-adrenalin-solution (10 mg/ml; Orion Pharma, Finland) was injected between the skull and the scalp for local anaesthesia and to prevent bleeding. Buprenorphine 0.1 mg/kg s.c. (Temgesic^®^ 0.3 mg/ml, Indivior UK Limited, United Kingdom) was administered before and 12 hours after the surgery for analgesia. After at least 4 days of recovery, a microdialysis probe (MAB 4.9.1.Cu; AgnTho’s AB) was inserted into the guide cannula and dialysis was started with Ringer solution (147 mM NaCl, 1.2 mM CaCl_2_, 2.7 mM KCl, 1.0 mM MgCl_2_, and 0.04 mM ascorbic acid) at a flow rate of 2 μl/min. Sample collection started after 2 hours of stabilization at 15 minute intervals. BT13 from 10 mM DMSO-stock was dissolved in Ringer solution until desirable concentration (the highest tested concentration (50 μM) was limited by BT13 solubility). The solutions were sonicated to enhance solubility of BT13. The final solutions contained less than 0.5% of DMSO, which alone did not significantly influence extracellular dopamine levels (data not shown). Concentration of dopamine was analyzed with high-performance liquid chromatography (HPLC) using electrochemical detection (Coulochem II; ESA, Inc., USA). The column (Kinetex 2.6u; XB-C18; 50 × 4.6 mm; Phenomenex; USA) was kept at 45°C with a column heater. The flow rate of mobile phase (0.1 M NaH_2_PO_4_, pH 4.0, 0.1 mg/ml octanesulphonic acid, 1.0 mM EDTA and 8% methanol) was 1 ml/min. An autoinjector (SIL-20AC, Shimadzu, Japan) was used to inject 25 μl of the sample into the chromatographic system. After achieving a stable baseline (determined as an average of 4 consecutive samples) with Ringer solution, BT13 was delivered into the striatum as a continuous infusion by reverse dialysis until the end of the experiment.

### Microinjections and stereotaxic surgery

Bilateral microinjections into the mouse dorsal striatum (AP = +0.6; ML = +/−1.8; DV = −2.2 relative to the bregma, according to the mouse brain atlas (Paxinos and Franklin, 2001) were performed using stereotaxic frame (Stoelting, USA). The stereotaxic surgery was conducted under isoflurane (Vetflurane 1000 mg/g, Virbac) anaesthesia. The animals received buprenorphine 0.1 mg/kg s.c. (Temgesic^®^ 0.3 mg/ml, Indivior UK Limited) for analgesia. A small amount of lidocaine-adrenalin-solution (10 mg/ml; Orion Pharma) was injected between the skull and the scalp for local anaesthesia and to prevent bleeding. Tested substances were delivered into the striata using an electronic injector (Quintessential stereotactic injector, Stoelting, USA) with a 10 l microsyringe (World Precision Instruments, United Kingdom). Injection speed was set to 0.2 μl/min and injection volume to 2 μl. Left striatum received BT13 or GDNF dissolved in saline with 0.5 % DMSO (vehicle). The right striatum was always injected with the vehicle. BT13 was injected in the doses of 103.5 μg (≈100 μg) N = 4), 207 μg (≈200 μg) (N = 4), 517.5 μg (≈500 μg) (N = 4) and 776.25 μg (≈750 μg) (N = 4) and GDNF in the dose of 5 or 10 μg (N = 4). At the completion of the injection, the needle was kept in place for 4 minutes and then slowly withdrawn to minimize backflow of the solution. The animals were allowed to recover in their home cage on a heating pad for 1 hour after the injection. Afterwards the mice were anesthetized with sodium pentobarbital (100 mg/kg, i.p.; Mebunat Vet 60 mg/ml, Orion Pharma) and transcardially perfused with warm PBS for 4 minutes followed by warm 4% PFA in 0.1 M phosphate buffer (pH 7.4) for 7 minutes. The brains were removed and postfixed in 4% PFA overnight at room temperature.

### 6-Hydroxydopamine lesion

Catecholaminergic neurotoxin 6-OHDA was injected unilaterally into the left dorsal striatum (AP = +1.0, ML = +2.7, DV = −4.0 relative to the bregma, according to the rat brain atlas (Paxinos and Watson, 1998) of male Wistar rats in a stereotaxic surgery under isoflurane (Vetflurane^®^ 1000 mg/g, Virbac) anaesthesia (Voutilainen *et al.*, 2009). Rats were fixed on a stereotaxic frame (Stoelting) using ear bars and incisor bar. Lidocaine-adrenalin-solution (10 mg/ml; Orion Pharma) was injected between the skull and the scalp for local anaesthesia and to prevent bleeding. The skull was exposed and a burr hole was made using a high-speed drill. An electronic injector (Quintessential stereotactic injector, Stoelting) and a 10 l microsyringe (Hamilton Company, USA) were used to deliver 16 μg of 6-OHDA (Sigma-Aldrich, Germany; calculated as free base and dissolved in ice-cold saline with 0.02% ascorbic acid). Injection rate was set to 1.0 μl/min and injection volume to 4 μl. At the completion of the injection, the needle was kept in place for 4 minutes and then slowly withdrawn. Desipramine (15mg/kg i.p.; calculated as free base; Sigma-Aldrich) was administrated 30 minutes before the 6-OHDA injection to prevent the uptake of 6-OHDA into noradrenergic and serotonergic nerve terminals.

### Administration of BT13 and GDNF

Another stereotaxic surgery was performed 1 hour after the 6-OHDA injection. The rats were anesthetized in the same manner as described above and the skull was exposed. A brain infusion cannula was implanted into the left dorsal striatum using the same coordinates as for the 6-OHDA injection (AP = +1.0; ML = +2.7; DV = −4.0) and secured to the skull with 3 stainless steel screws and dental cement (Aqualox). The cannula was connected via a 3 cm-long catheter tubing to an osmotic infusion pump (Alzet model 2002, Durect Co., USA) which was placed into a subcutaneous pocket between scapulae. The pumps constantly infused BT13 (0.25-0.5 μg/μl in 100% propylene glycol), GDNF (0.25 μg/μl in PBS) or 100% propylene glycol (vehicle-treated group) into the striatum at a flow rate of ∼0.5 μl/h for 7 days. The resulting dose for BT13 was ∼3-6 μg/24 h and for GDNF ∼3 μg/24 h. The dose for BT13 could not be determined accurately because the solubility of BT13 in 100% propylene glycol was limited to 0.5 μg/μl which was the highest concentration that remained dissolved for 7 days in stabile conditions in a test tube at 37°C. However, changes in ambient conditions during the pump implantation and 7-day infusion period may have precipitated some of the compound. In addition, variation in the mean pumping rate between different batches of the osmotic pumps may have resulted in dose deviation of more than 1 μg/24 h at the concentration of 0.5 μg/μl. At the end of the infusion, 7 days post lesion, the pump, cannula, dental cement and screws were removed in the third stereotaxic surgery under isoflurane anaesthesia and the incision was sutured.

Before the stereotaxic surgeries rats received buprenorphine 0.05 mg/kg s.c. (Temgesic^®^ 0.3 mg/ml, Indivior UK Limited) for analgesia. Carprofen 5 mg/kg (Rimadyl Vet^®^ 50mg/ml, Zoetis Inc., NJ, USA) was injected s.c. immediately after the surgeries to relieve postoperative pain. Additional doses of buprenorphine and carprofen were given 1 day after the surgeries.

### Behavioral analysis

D-Amphetamine-induced rotational behavior was measured 2, 4 and 6 weeks after the 6-OHDA lesion in automatic rotometer bowls (Med Associates Inc., VT, USA) as described previously (Ungerstedt and Arbuthnott, 1970; Lindholm *et al.*, 2007). After a habituation period of 30 minutes, rats received a single injection of D-amphetamine (2.5mg/kg i.p.; calculated as free base; Division of Pharmaceutical Chemistry and Technology, University of Helsinki, Finland). Automated rotation sensors recorded full (360°) uninterrupted turns for a period of 120 minutes. Net ipsilateral turns to the lesion side were calculated by subtracting contralateral turns from ipsilateral turns. After completion of the last behavioral test, rats were anesthetized with sodium pentobarbital (90 mg/kg, i.p.; Mebunat Vet^®^ 60 mg/ml, Orion Pharma) and perfused transcardially first with warm PBS for 5 minutes followed by warm 4% PFA in PBS for 10 minutes. The brains were excised, post-fixed in 4% PFA overnight at 4°C and stored in 20% sucrose in PBS at 4°C until freezing in dry ice-cooled isopentane.

### Immunohistochemistry

Immunohistochemical staining of pERK1/2 and pS6 were performed on 5 μm thick coronal sections of paraffin-embedded mouse brains using primary antibodies raised against pERK1/2 (1:300, Cell Signaling Technology Cat# 4370, RRID:AB_2315112) and pS6 (1:300, Cell Signaling Technology Cat# 5364, RRID:AB_10694233). To visualize the stainings, sections were incubated with HRP-conjugated goat anti-rabbit secondary antibody (1:500, Sigma-Aldrich Cat# A6154, RRID:AB_258284), washed 3 times in TBS and treated with 3,3’-diaminobenzidine (DAB; Cat# SK-4100, USA) as a chromogen.

To assess the number of remaining dopamine neurons in the SNpc and the density of dopaminergic fibers in the striatum, rat brains were cut into 40-μm thick free-floating coronal cryosections. For dopamine transporter (DAT) staining antigen retrieval was performed by incubating the sections in 10 mM citrate buffer (pH 6) for 30 minutes at 80°C. The sections were probed with monoclonal mouse-anti-TH antibody (1:2000, Millipore Cat# MAB318, RRID:AB_2201528) or with monoclonal rat-anti-DAT antibody (1:2000, Millipore Cat# MAB369, RRID:AB_2190413) as described previously (Voutilainen et al., 2009; Bäck et al., 2013). After rinsing in PBS, for TH staining the sections were incubated for 2 hours at room temperature in biotinylated horse-anti-mouse secondary antibody solution (1:200, Vector Laboratories Cat# BA-2001, RRID:AB_2336180), and for DAT staining for 1 hour at room temperature in biotinylated rabbit-anti-rat secondary antibody solution (1:200, Vector Laboratories Cat# BA-4000, RRID:AB_2336206). The stainings were reinforced with avidin-biotinylated HRP complex (Vectastain Elite ABC HRP Kit, Vector Laboratories Cat# PK-6100, RRID:AB_2336819), rinsed with PBS and visualized using DAB (0.5 mg/ml in 0.03% H_2_O_2_ in PBS; Cat# SK-4100, USA) as a chromogen.

Stained sections were dehydrated in series of ethanol solutions with increasing concentrations, clarified in xylen and finally mounted in DePeX^®^ mounting medium (VWR International Ltd., England). In all cases endogeneous peroxidase activity was quenched by preincubation of sections with 3% H_2_O_2_ solution.

### Analysis of pERK and pS6 immunohistochemistry

The stained sections were scanned using an automated bright field microscopy slide scanner (3DHistech Ltd., Hungary). Mean optical density (OD) of the staining in a single section (closest to the injection site and with the highest signal) from both hemispheres of each animal was analyzed using ImageJ software (Media Cybernetics Inc, USA). In cases where the highest signal was not located in the same section for both hemispheres, the values were counted from separate sections. Digital images of the sections were first converted into 8-bit grey scale and inverted. Then roughly equal sized areas were outlined from both hemispheres and OD values were counted within these areas. Background OD values were measured from the peripheral striatal area that lacked the signal or from the septum and subtracted from the OD values in the areas selected for analysis. Resulting data were normalized to the area of analysed selections and subjected to the statistical analysis. For presentation purpose the values for BT13- and GDNF-treated sides were normalized to the values for vehicle-treated side of the same brain.

### Stereological assessment of TH-positive cells in the SNpc

TH-positive cell bodies in the SNpc of rat brain were counted by a person blinded to the treatment groups using Olympus BX51 (Olympus Corporation, Japan) microscope connected to a computer running Stereo Investigator software version 11.06.2 (MBF Bioscience, USA) as described previously (Lindholm *et al.*, 2007; Voutilainen *et al.*, 2009). Optical fractionator probe with systematic random sampling was used for cell counting in Stereo Investigator and the parameters were set to the following values in each case: grid size: 92×92 m; counting frame size: 80×80 m; dissector height: 13 m; guard zones: 1.5 m at the bottom and top of section.

SNpc was outlined with a contour under 4× magnification (Olympus UPlanFl 4×/0.13 objective) and the subsequent cell counting was done under 60× magnification (Olympus PlanApo 60× 1.40 Oil /0.17 objective) using Sigma-Aldrich immersion oil (Cat# 56822). TH-positive cell bodies were bilaterally counted in the SNpc on 3 coronal sections at approximately the same rostro-caudal sites, which were visually identified by the following landmarks: (1) medial terminal nucleus of the accessory optic tract (MT) partially divides the SNpc, (2) MT divides the SNpc into two parts, and (3) clusters of blood vessels enter the midbrain from the ventral side towards the SN without MT being present on that section. The counting contour always excluded the lateral part of the SNpc. In the case of (1) and (2) the medial edge of the SNpc was limited by the MT. In the case of (3), approximately equal distance from the lateral edge of the SNpc was taken to the medial direction on both sides with the aim to count only the SNpc and to exclude counting of the ventral tegmental area cells. Cells needed to have at least one long neurite and be polygonal to be included in the counting. The results are expressed as mean number of TH-positive cells per section from the 3 sections.

### TH-positive fiber density in the striatum

Analysis of TH-positive fibers in the striatum was performed under blinded conditions. Optical density of TH-positive fibers was measured bilaterally from 3 coronal sections of each rat at approximately the same rostro-caudal sites (AP = +1.2; +0.48 and –0.26 relative to the bregma, according to the rat brain atlas (Paxinos and Watson, 1998). Digital images of the TH-immunostained sections were acquired with an automated bright field microscopy slide scanner (3DHistech Ltd.). Obtained images were converted to 8-bit grey scale. After inverting the colours, integrated optical densities of the dorsal striata divided by the measured area were analyzed with Fiji ImageJ software (Media Cybernetics Inc, USA). Nonspecific background staining was measured from corpus callosum and subtracted from the striatal optical densities. The data are presented as percentage of the lesioned striatum as compared with the intact striatum.

